# Controlling Noise in the Timing of Intracellular Events: A First-Passage Time Approach

**DOI:** 10.1101/056945

**Authors:** Khem Raj Ghusinga, John J. Dennehy, Abhyudai Singh

## Abstract

In the noisy cellular environment, gene products are subject to inherent random fluctuations in copy numbers over time. How cells ensure precision in the timing of key intracellular events, in spite of such stochasticity is an intriguing fundamental problem. We formulate event timing as a first-passage time problem, where an event is triggered when the level of a protein crosses a critical threshold for the first time. Novel analytical calculations are preformed for the first-passage time distribution in stochastic models of gene expression, including models with feedback regulation. Derivation of these formulas motivates an interesting question: is there an optimal feedback strategy to regulate the synthesis of a protein to ensure that an event will occur at a precise time, while minimizing deviations or noise about the mean. Counter-intuitively, results show that for a stable long-lived protein, the optimal strategy is to express the protein at a constant rate without any feedback regulation, and any form of feedback (positive, negative or any combination of them) will always amplify noise in event timing. In contrast, a positive feedback mechanism provides the highest precision in timing for an unstable protein. These theoretical results explain recent experimental observations of single-cell lysis times in bacteriophage *λ*. Here, lysis of an infected bacterial cell is orchestrated by the expression and accumulation of a stable *λ* protein up to a threshold, and precision in timing is achieved via feedforward, rather than feedback control. Our results have broad implications for diverse cellular processes that rely on precise temporal triggering of events.

## 1. INTRODUCTION

Timing of events in many cellular processes, such as cell-cycle control [1–4], cell differentiation [5, 6], sporulation [7, 8], apoptosis [9–11], development[12, 13], temporal order of gene expression [14–16], depend on regulatory proteins reaching critical threshold levels. Triggering of these events in single cells are influenced by fluctuations in protein levels that arise naturally due to noise in gene expression [17–26]. Increasing evidence shows considerable cell-to-cell variation in timing of intracellular events among isogenic cells [27–29], and it is unclear how noisy expression generates this variation. Characterization of controls strategies that buffer stochasticity in event timing are critically needed to understand reliable functioning of diverse intracellular pathways that rely on precision in timing.

Mathematically, noise in the timing of events can be investigated via the first-passage time (*FPT*) framework, where an event is triggered when a stochastic process (single-cell protein level) crosses a critical threshold for the first time. There is already a rich tradition of using such first-passage time approaches to study timing of events in biological and physical sciences [30–43]. Following this tradition, exact analytical expression for the *FPT* distribution are computed in experimentally validated and commonly used stochastic models of gene expression. These results provide novel insights into how expression parameters shape statistical fluctuations in event timing.

To investigate control mechanisms for buffering noise in timing, we consider feedback regulation in protein synthesis, where the expression rate varies arbitrarily with the protein count. Such feedback can be implemented directly through auto-regulation of gene promoter activity by its own protein [44–47] or indirectly via intermediate states [48]. It is important to point out that while the effects of such feedback loops on fluctuations in protein copy number are well studied [49–53], their impacts on stochasticity in event timing have been overlooked. We specifically formulate the problem of controlling precision in event timing as follows: what optimal form of feedback regulation ensures a given mean time to an event, while minimizing deviations or noise about the mean. It turns out that for certain minimal models of stochastic expression this optimization problem can be solved analytically, providing counter-intuitive insights. For example, in many cases negative feedback regulation is found to amplify noise in event timing, and in some cases, the optimal form of feedback is to not have any feedback at all. The robustness of these result are analyzed in the context of different noise mechanisms, such as intrinsic versus extrinsic noise in gene expression [54–57]. Finally, we discuss in detail how our results explain recent experimental observations of single-cell lysis times in bacteriophage *λ*, where precision in timing is obtained without any feedback regulation.

**FIG. 1.**
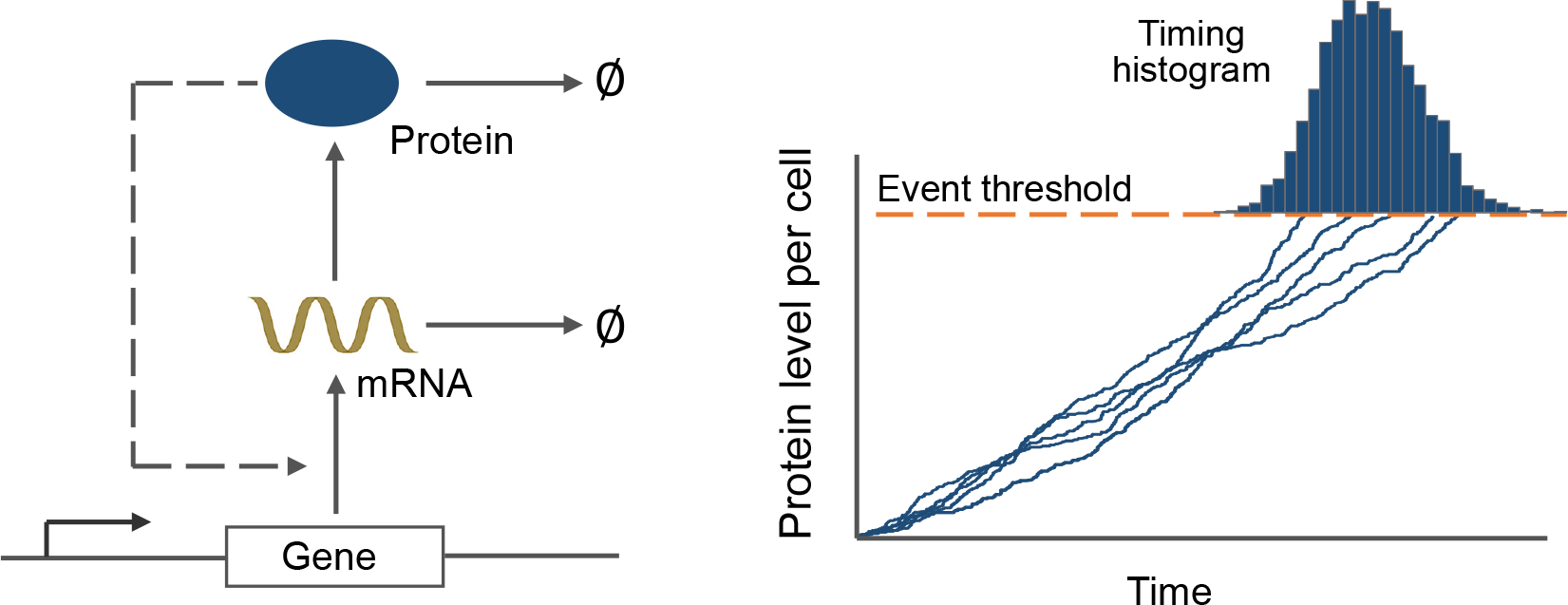
Modeling event timing as a first-passage time problem. *Left*: Model schematic with a gene transcribing mRNAs, which is further translated into proteins. The rate of transcription is assumed to be regulated by the protein level, creating a feedback loop. *Right*: The timing of an intracellular event is formulated as the first-passage time for the protein level to reach a critical threshold. Sample trajectories for the protein level over time obtained via Monte Carlo simulations are shown, and they cross the threshold at different times due to stochasticity in gene expression. The histogram of the event timing based on 5,000 trajectories is shown on the top.

## II. STOCHASTIC MODEL FORMULATION

Consider a gene that is switched on at time *t* = 0 and begins to express a timekeeper protein. The intracellular event of interest is triggered once the protein reaches a critical level in the cell. We describe a minimal model of protein synthesis that incorporates two key features - expression in random bursts and feedback regulation (Fig. 1). Let *x*(*t*) ∈ {0,1,…} denote the level of a protein in a single cell at time *t*. When *x*(*t*) = *i*, the gene is transcribed at a Poisson rate *k_i_*. Any arbitrary form of feedback can be realized by appropriately defining *k_i_* as a function of *i*. For example, increasing (decreasing) *k_i_*’s correspond to a positive (negative) feedback loop in protein production, and a fixed transcription rate implies no feedback. Assuming short-lived mRNAs, each mRNA degrades instantaneously after producing a burst of *B* protein molecules [58–62]. In agreement with experimental and theoretical studies [63–65], *B* is assumed to follow a geometric distribution

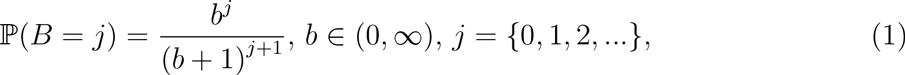

where *b* denotes the mean protein burst size and the symbol 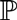 is the notion for probability. Finally, each protein molecule degrades with a constant rate *γ*. The time evolution of *x*(*t*) is described through the following probabilities of occurrences of burst and decay events in the next infinitesimal time (*t*, *t* + *dt*]

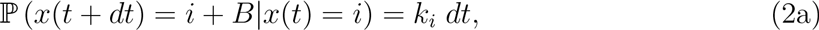

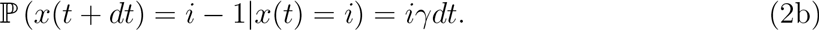

Note that in this representation of gene expression as a bursty birth-death process, the mRNA transcription rate *k_i_* is the burst arrival rate, while the rate at which proteins are translated from an mRNA determines the mean protein burst size *b*. Next, we formulate event timing through the first-passage time framework.

## III. COMPUTING EVENT TIMING DISTRIBUTION

The time to an event is the first-passage time for *x*(*t*) to reach a threshold *X* starting from a zero initial condition *x*(0) = 0 (Fig. 1). It is mathematically described by the following random variable

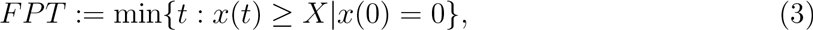

and can be interpreted as the time taken by a random walker to first reach a defined point. Our goal is to obtain closed-form expressions for the *FPT* statistics in terms of underlying model parameters. Note that if the protein did not decay, then *x*(*t*) accumulates over time and the *FPT* distribution is obtained by observing

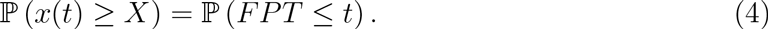

However, with protein degradation, the *FPT* calculation needs careful consideration so as to avoid counting multiple crossings of the threshold.

To compute the *FPT* we imagine the bursty birth-death process on a finite state-space [0,1,…, *X*], where the states represent the protein count (Fig. 2). All states denoting *x*(*t*) ≥ *X* are combined into a single absorbing state *X*. In this model, the probability of the protein level reaching *X* in the small time window (*t*,*t* + *dt*) is the probability of being in state *i* at time *t*, and a jump of size *X* − *i* or larger occurs in (*t*,*t* + *dt*). Using the fact that for a geometrically distributed burst size *B*,

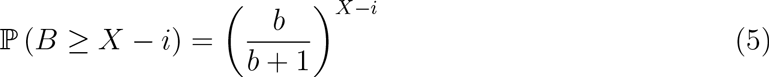

and rate of burst arrival is *k_i_* when *x*(*t*) = *i*, the probability density function (pdf) of the first-passage time is giving by

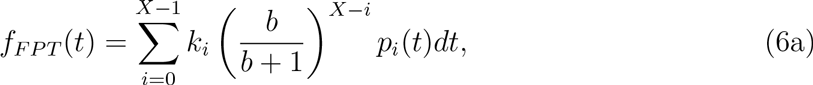

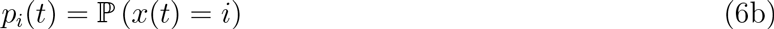

**FIG. 2.**
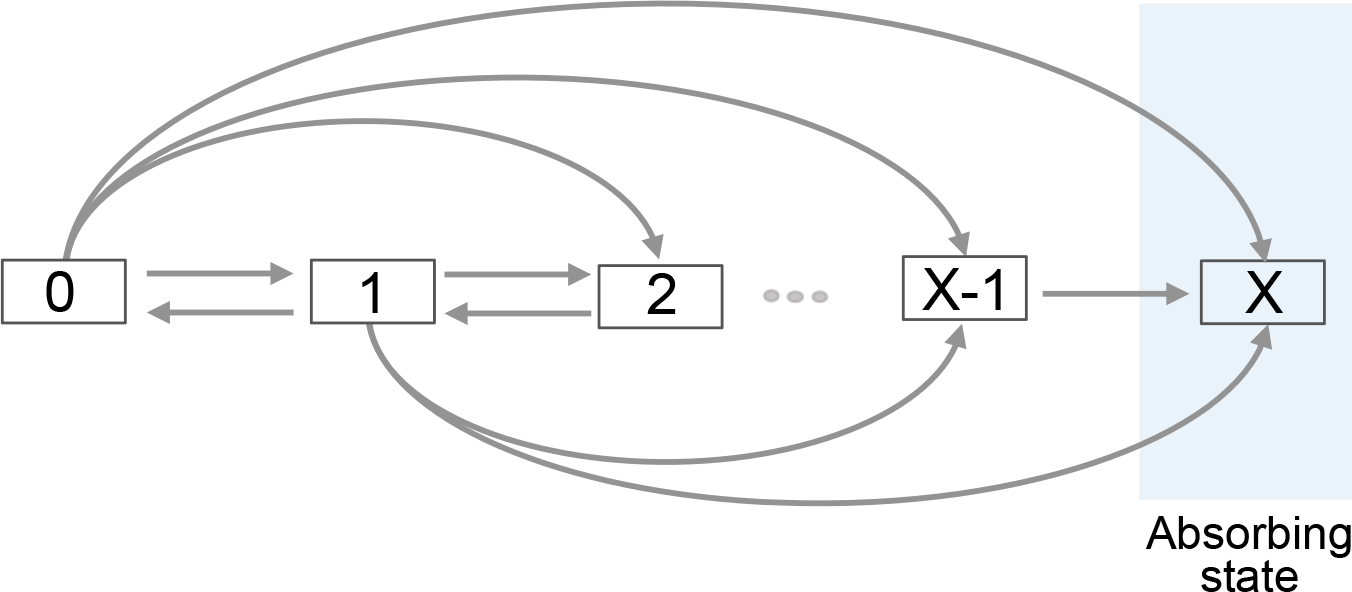
Illustration of a bursty birth-death process for computing the first-passage time. States [0,1,…, *X*] represent the protein population counts, and arrows represent transition between states due to burst and decay events. The destination of a forward jump (a birth event) is decided by the burst size while each degradation event reduces the protein count by one. The process terminates when the protein level reaches the absorbing-state *X* and the first-passage time is recorded.

The pdf can be compactly written as a product

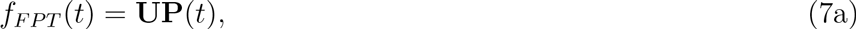

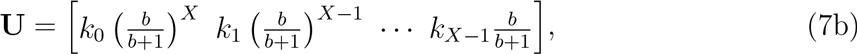

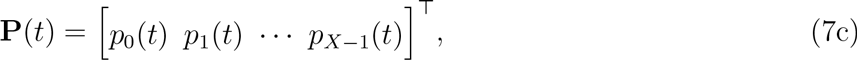

where **U** is a row vector of 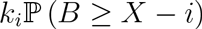 and **P**(*t*) is a column vector of *p_i_*(*t*). The time evolution of **P**(*t*) is given by the linear dynamical system

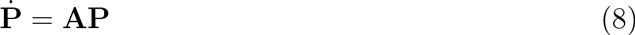

derived from the Chemical Master Equations (CME) corresponding to the bursty birth-death process [66, 67]. It turns out that, in this case the matrix **A** is a Hessenberg matrix whose *i^th^* row and *j*^th^ column element is given by

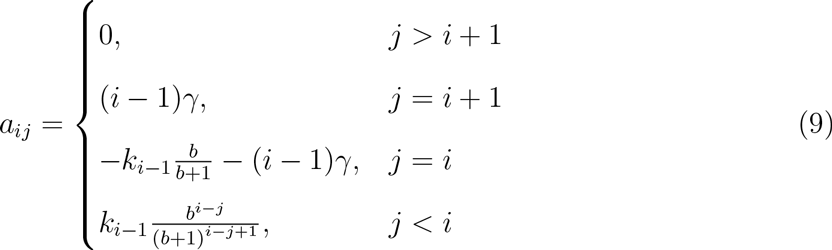

*i*,*j* ∈ {1,…, *X*}. Solving (8) and using (7a) yields the following pdf for the first-passage time

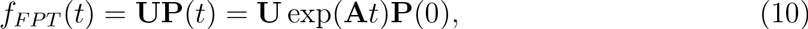

where **P**(0) = [1 0… 0]^*T*^ is vector of probabilities at *t* = 0 that follows from *x*(0) = 0. While this pdf provides complete characterization of the event timing, we are particularly interested in the lower-order statistical moments of *FPT*. Next, we exploit the structure of the **A** matrix to obtain analytical formulas for the first and second order moments of the first-passage time.

## IV. MOMENTS OF THE FIRST-PASSAGE TIME

From (10), the *m^th^* order uncentered moment of the first-passage time is given by

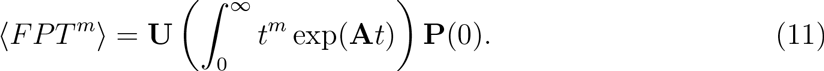

Since the matrix **A** is full-rank with negative eigenvalues (see section S1 of the SI), the above integral can be computed as

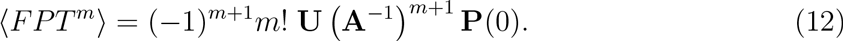

Using the inverse of a Hessenberg matrix, we obtain the following first two moments of *FPT* (see section S2 of the SI)

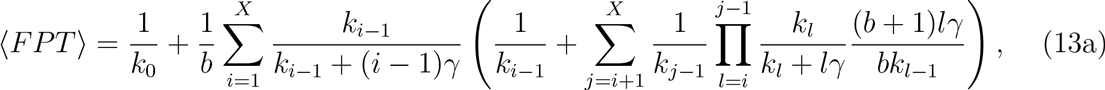

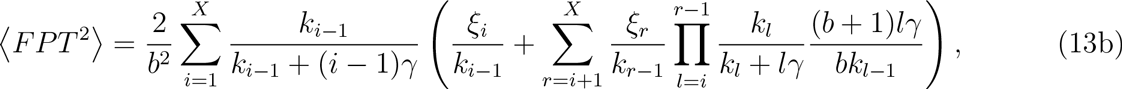

where *δ*_*i*−1_ represents the Kronecker delta which is one if *i* =1 and zero otherwise, and

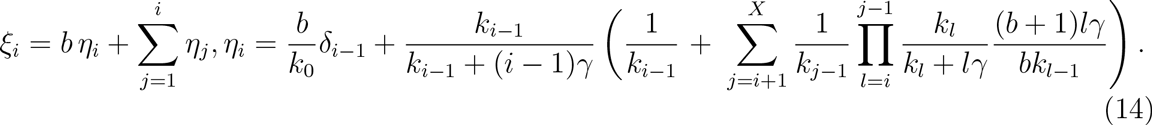

These results represent the first analytical computations of the *FPT* statistics for a bursty-birth death process with a random burst size and a state-dependent burst arrival rate (i.e., feedback regulation in transcription).

We investigate the complex formulas in (13) for some limiting cases. For a stable long-lived protein (*γ* = 0) and a constant transcription rate (no feedback; *k_i_* = *k*), moment expressions reduce to

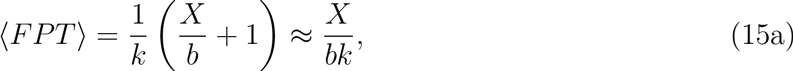

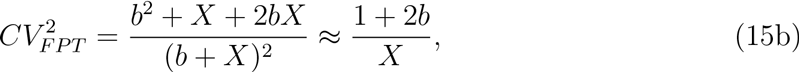

where 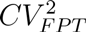 represent the noise in the first-passage time as quantified by its coefficient of variation squared. The approximate formulas in (15) are valid for a high event threshold compared to the mean protein burst size (*X*/*b* ≫ 1). These formulas reveal important insights, such as, the noise in **FPT** is invariant of the transcription rate *k*. Moreover, ⟨*FPT*⟩ and 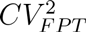 can be independently tuned — increasing the event threshold and/or reducing the burst size will lower the noise level. Once 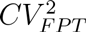 is sufficient reduced, *k* can be modulated to adjust the mean event timing to any desired value. Next, we explore how feedback regulation of the transcription rate impacts noise in timing, for a given *X* and *b*.

## V. OPTIMAL FEEDBACK STRATEGY

Having derived the *FPT* moments, we investigate optimal forms of transcriptional feedback that schedule an event at a given time with the highest precision. Mathematically, this corresponds to an constraint optimization problem: find transcription rates *k*_0_, *k*_1_,…, *k*_*X*−1_ that minimize ⟨*FPT*^2^⟩ for a fixed ⟨*FPT*⟩. We first consider a stable protein whose half-life is much longer than the event timescale, and hence, degradation can be ignored (*γ* = 0).

### A. Optimal feedback for a stable protein

When the protein of interest does not decay (*γ* = 0), the expressions for the *FPT* moments take much simpler forms

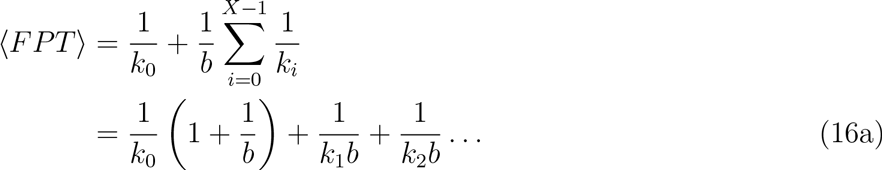

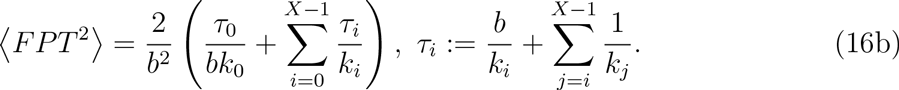

Note that in (16a) the contribution of *k*_0_ (transcription rate when there is no protein) is quite different from the other transcription rates *k_i_*, *i* ∈ {1, 2,…, *X* − 1}. For instance, when the event threshold is large compared to the mean burst size (*X* ≫ *b*), then the term 1/*k*_0_ can be ignored and 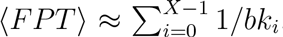. In contrast, if the burst size is large (*b* ≪ *X*) then ⟨*FPT*⟩ ≈ 1/*k*_0_, as a single burst event starting from zero protein molecules is sufficient for threshold crossing. Similar observation for different contributions of *k*_0_ can be made about (16b).

It turns out that, for these simplified formulas, the problem of minimizing ⟨*FPT*^2^⟩ given ⟨*FPT*⟩ can be solved analytically using the method of Lagrange multipliers (see section S3 of the SI for details). The optimal transcription rates are given by

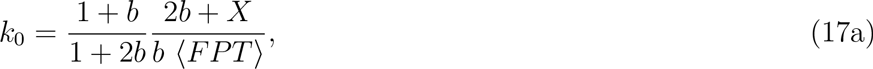

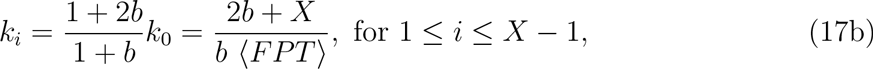

and all rates are equal to each other except for *k*_0_. Intuitively, the difference for *k*_0_ comes from the fact that it contributes differently to the *FPT* moments as compared to other rates. Note that for a small mean burst size (*b* ≪ 1), *k*_0_ = *k_i_*, whereas *k*_0_ = *k_i_*/2 for a sufficiently large *b*. Despite this slight deviation in *k*_0_, for the purposes of practical implementation, the optimal feedback strategy in this case is to have a constant transcription rate (i.e., no feedback in protein expression).

**FIG. 3.**
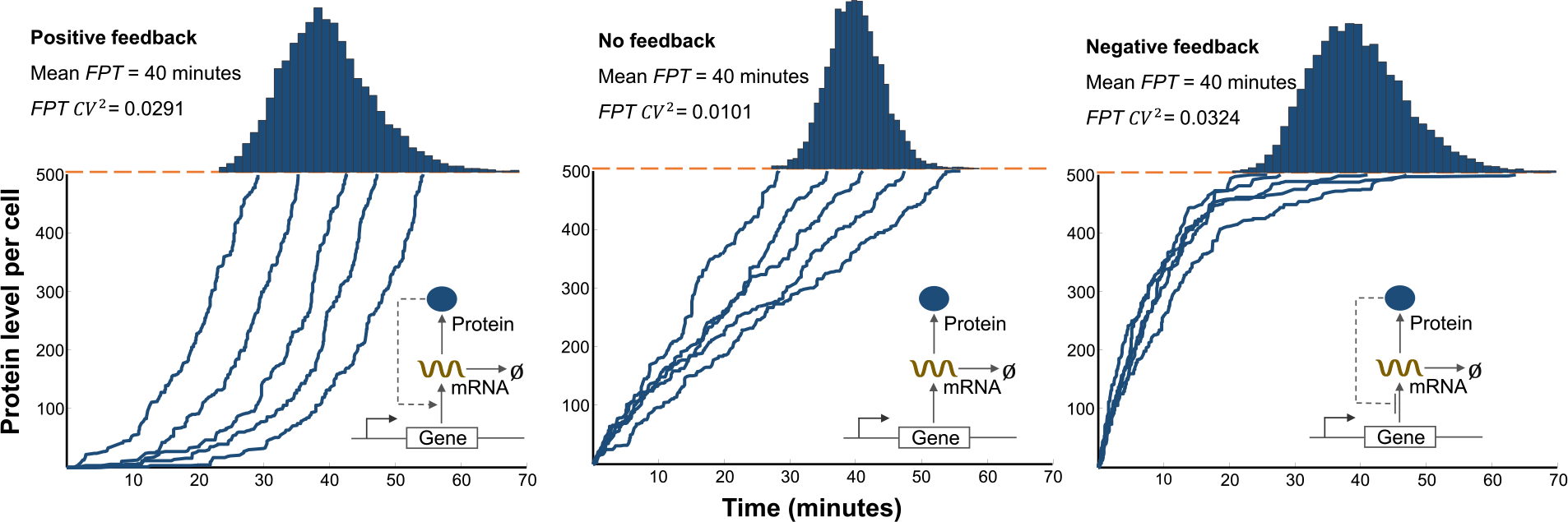
For a stable protein, no feedback provides the lowest noise in event timing for a fixed mean *FPT*. Protein trajectories obtained using the Stochastic Simulation Algorithm (SSA) for a stochastic gene expression model with positive feedback (left), no feedback (middle), negative feedback (right) [68]. The threshold for event timing is assumed to be 500 protein molecules and feedback is implement by assuming a transcription rate of the form *k_i_* = *c*_1_ ± *c*_2_*i*, where *c*_2_ = 0 (no feedback), *c*_2_ = 0.05min^−1^ (positive feedback), *c*_2_ = −0.05min^−1^ (negative feedback). For a given value of *c*_2_, the mean *FPT* is kept constant (40 minutes) by changing the parameter *c*_1_. The mRNA half-life is assumed to be 2.7 min, and proteins are translated from mRNAs at a rate 0.5 min^−1^, which corresponds to a mean burst size of *b* = 2. Histograms on the top represent distribution of *FPT* from 10,000 Monte Carlo simulations.

We illustrate the above result via Monte Carlo simulations of stochastic gene expression models that explicitly include mRNA dynamics (Fig. 3). Here, feedback is implemented using linear transcription rates

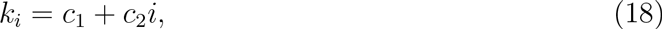

where *c*_2_ = 0 (no feedback), *c*_2_ > 0 (positive feedback), *c*_2_ < 0 (negative feedback), and |*c*_2_| is referred to as the feedback strength. As expected from theory, a no feedback strategy outperforms negative/positive feedbacks in terms of minimizing deviations in *FPT* around a given mean event time (Fig. 3). While similar results were obtained for implementing transcription rates using Hill functions (see section S4 of the SI), we prefer to use linearized rates as they have a fewer number of parameter and a clearer notion of feedback strength.

### B. Optimal feedback for an unstable protein

Now consider the scenario where protein degradation cannot be ignored over the event timescale (*γ* ≠ 0). Unfortunately, the expressions of the *FPT* moments in (13) are too convoluted for the optimization problem to be solved analytically, and the effect of different feedbacks is investigated numerically. Our strategy is as follows: choose a certain feedback strength *c*_2_ in (18), appropriately tune *c_1_* using (13b) for the desired mean event timing, and explore the corresponding noise in *FPT* as measured by its coefficient of variation squared 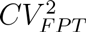 (additional details in section S5 of the SI). Counter-intuitively, results show that for a given value of *γ*, a negative feedback loop in gene expression has the highest 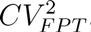, and its performance deteriorates with increasing feedback strength (Fig. 4A). In contrast, 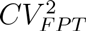 first decreases with increasing strength of the positive feedback, and then increases after an optimal feedback strength is crossed (Fig. 4A). Thus, when the protein is not stable, precision in timing is attained by having a positive feedback in protein synthesis with an intermediate strength.

We next explore how the minimal achievable noise in event timing, for a fixed ⟨*FPT*⟩, varies with the protein decay rate *γ*. Our analysis shows that the minimum 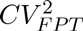 obtained via positive feedback increases monotonically with *γ*, and 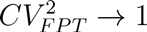 as *γ* ⟶ ∞ (Fig. 4B). Recall that the coefficient of variation of an exponentially distributed random variable is exactly equal to one. Thus, as the protein becomes more and more unstable, the timing process becomes memoryless yielding exponentially distributed first-passage times. A few interesting observations can be made from Fig. 4B: i) Higher protein burst sizes result in much larger 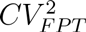 and a faster approach to 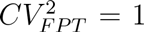 as *γ* ⟶ ∞; ii) The difference in 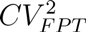 for optimal feedback and no feedback is indistinguishable when the protein is stable (*γ* = 0) or highly unstable (*γ* ⟶ ∞); and iii) For a range of intermediate protein half-lives, the optimal feedback strategy provides an order of magnitude better suppression of 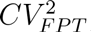, as compared to no feedback regulation. Taken together, these observations suggest that the timescale of protein turnover and the extent of bursty expression sets a fundamental limit to how much statistical fluctuations in *FPT* can be buffered.

Consistent with our above analysis, we find that a positive feedback mechanism provide the highest precision in timing in more complex stochastic expression models that explicitly include mRNA dynamics (Fig. 4C). An interesting point to note is that the protein trajectories for the optimal positive feedback case look fairly linear, and similar to the trajectories seen in the no feedback case when *γ* = 0 (compare rightmost plot in Fig. 4C with middle plot in Fig. 3). One way to think about this is to consider protein synthesis in the deterministic limit described by the following ordinary differential equation

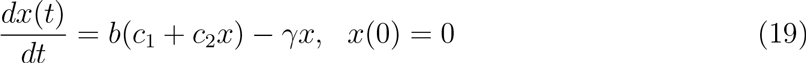

with mean protein burst size *b* and a linear feedback form (18). If the feedback strength is chosen as

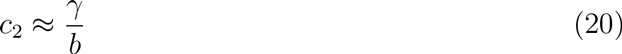

then the time evolution of *x*(*t*) would be linear over time in (19). Indeed, our detailed stochastic analysis shows that the optimal feedback strength that minimizes 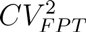 in the stochastic model is qualitatively similar but not identical to (20) (see section S5 of the SI).

**FIG. 4.**
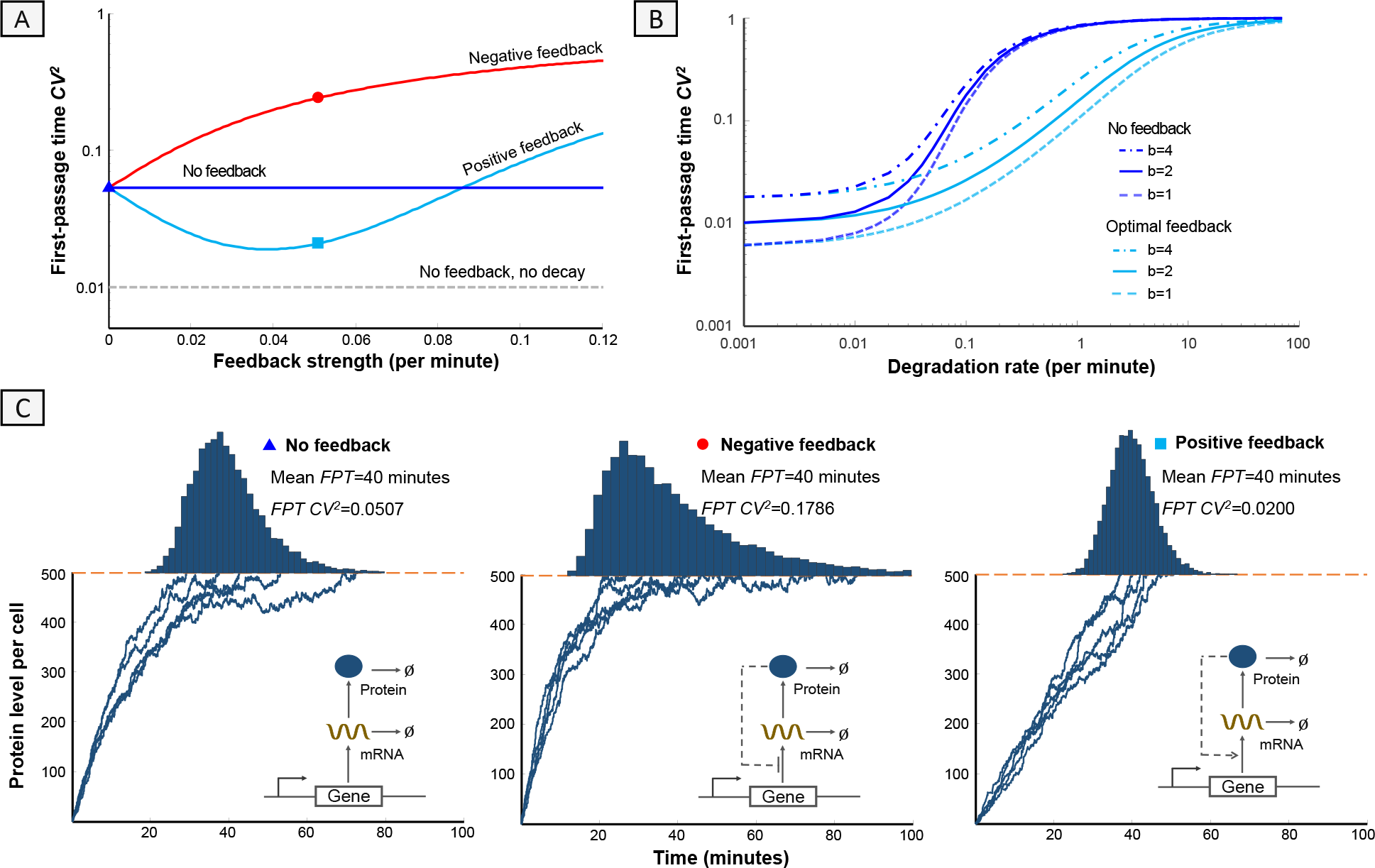
For an unstable protein, positive feedback provides the lowest noise in event timing for a fixed mean *FPT*. (A) Noise in timing 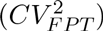 as a function of the feedback strength |*c*_2_| for different control strategies. The value of *c*_1_ is changed in (18) so as to keep ⟨*FPT*⟩ = 40 mins fixed. The performance of the negative feedback worsens with increasing feedback strength. In contrast, positive feedback with an optimal value of *c*_2_ provides the highest precision in event timing. Other parameters used are *γ* = 0.05 min^−1^, *X* = 500 molecules, *b* = 2. The optimal value attained via positive feedback is much higher than the minimal value of 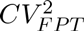 for a stable protein (dashed line). (B) The minimum 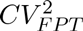 obtained via positive feedback increases monotonically with the protein degradation rate. For comparison purposes, 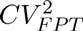 obtained without any feedback (*c*_2_ = 0) is also plotted. (C) Monte Carlo simulations of protein trajectories under different forms of feedback control. A positive feedback outperforms the other feedback strategies in terms of minimizing statistical fluctuations in event timing. Histograms of the first-passage times obtained from 10, 000 Monte Carlo runs are shown on top. For these simulations *X* = 500 molecules, *γ* = 0.05 min^−1^, mRNA half-life is assumed to be 2.7 min, proteins are translated from mRNAs at a rate 0.5min^−1^, and *c*_2_ = 0 (no feedback), *c*_2_ = 0.05min^−1^ (positive feedback), *c*_2_ = −0.05 min^−1^ (negative feedback).

## VI. DISCUSSION

We have systematically investigated ingredients essential for precision in timing of biochemical events at the level of single cells. Our approach relies on modeling even timing as the first-passage time for a stochastically expressed protein to cross a threshold level. This framework was used to uncover optimal strategies for synthesizing the protein that ensures a given mean time to event triggering (threshold crossing), with minimal fluctuations around the mean. The main contributions and insights can be summarized as:

1. Novel analytical calculations for the first-passage time in stochastic models of gene expression, with and without feedback regulation.
2. If the protein half-life is much longer than the timescale of the event, the highest precision in event timing is attained by having no feedback, i.e., express the protein at a constant rate (Fig. 3).
3. In the absence of feedback, the noise in event timing is given by (15) and determined by the molecular threshold (*X*) and the protein burst size (*b*). Once *X* and *b* are chosen for a tolerable noise level, the mean time to the event can be adjusted independently through the transcription rate *k*.
4. If the protein half-life is comparable or shorter than the timescale of the event, then positive feedback provides the lowest noise in event timing (Fig. 4A). Moreover, negative feedback always amplifies noise around the mean time.
5. The minimum achievable noise in timing increases with the protein decay rate 7 and approaches 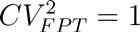 as *γ* ⟶ ∞ (Fig. 4B).

How robust are these findings to alternative noise sources and key modeling assumptions? For example, the model only considers noise from low-copy number fluctuation in gene product levels, and ignores any form of “extrinsic noise” that arises from cell-to-cell differences in gene expression machinery [55, 69]. To incorporate such extrinsic noise, we alter the transcription rate to *k_i_Z*, where *Z* is drawn from an a priori probability distribution at the start of gene expression (*t* = 0), and remains fixed till the threshold is reached. Interestingly, the optimal feedback derived in (17) does not change even after adding extrinsic noise to the transcription rate or the protein burst size (see section S6 of the SI). Another important model feature is geometrically distributed protein burst sizes, which follows from the assumption of exponentially distributed mRNA lifetimes. We have also explored the scenario of perfect memory in the mRNA degradation process, which results in a mRNA lifetime distribution given by the delta function. In this case, the protein burst size is Poisson and the optimal feedback strategy is fairly close to having no feedback for a stable protein (see section S7 of the SI). Next, we discuss the biological implications of our findings in the context of phage *λ*’s lysis times, i.e., the time taken by the virus to destroy infected bacterial cells.

### A. Connecting theoretical insights to *λ* lysis times

Phage *λ* has recently emerged as a simple model system for studying event timing at the level of single cells [27, 28]. After infecting *E. coli*, *λ* expresses a protein, holin, which accumulates in the inner membrane. When holin reaches a critical threshold concentration, it undergoes a structural transformation, forming holes in the membrane [70]. These holes allow other lysis proteins (endolysin and spanin) to access and rupture the cell wall [71]. Subsequently the cell lysis and phage progeny are released into the surrounding medium. Since hole formation and cell rupture are nearly simultaneous, lysis timing depends on denovo expression and accumulation of holin in the cell membrane up to a critical threshold [70–72]. Data reveals precision in the timing of lysis - individual cells infected by a single virus lyse on average at 65 mins, with a standard deviation of 3.5 mins, implying a coefficient of variation of ≈ 5%. Such precision is expected given the existence of an optimal lysis time [73–77]. Intuitively, if *λ* lysis is early then there are no viral progeny. In contrast, if *λ* lysis is late then the infected cell could die before lysis is effected, trapping the virus with it.

The threshold for lysis is reported to be a few thousand holin molecules [78]. Moreover, the holin mean burst size (average number of holins produced in a single mRNA lifetime) is estimated as *b* ≈ 1 − 3 [78]. Based on our *FPT* moment calculations in (15), such a small protein burst size relative to the event threshold will yield a tight distribution of lysis times. Interestingly, (15) provides insights for engineering mutant *λ* that lyse, on average, at the same time as the wild type, but with much higher noise. This could be done by lowering the threshold for lysis through mutations in the holin amino acid sequence [28], and also reducing the holin mRNA transcription or translation rate so as to keep the same mean lysis time. It is important to point out that since holin proteins are long-lived and do not degrade over relevant timescales [79], *λ*’s lysis system with no known feedback in holin expression provides better suppression of lysis-time fluctuation compared to any feedback regulated system.

### B. Additional mechanism for noise buffering

The surprising ineffectiveness of feedback control motivates the need for other mechanisms to buffer noise in event timing. Intriguingly, *λ* uses feedforward control to regulate the timing of lysis. Feedforward control is implemented through two proteins with opposing functions: holin and antiholin [80–82]. In the wild-type virus both proteins are expressed in a 2:1 ratio (for every two holins there is one antiholin) from the same mRNA through a dual start motif. Antiholin binds to holin and prevents holin from participating in hole formation, creating an incoherent feedforward circuit (Fig. 5). Synthesis of antiholin leads to a lower burst size for active holin molecules, and increases the threshold for the total number of holins needed for lysis - both factors functioning to lower the noise in event timing. Consistent with this prediction, variants of *λ* lacking antiholin are experimentally observed to exhibit much higher intercellular variation in lysis times as compared to the wild-type virus [28, 83]. In summary, *λ* encodes a multitude of regulatory mechanisms (low holin burst size; no feedback regulation; feedforward control) to ensure that single infected cells lyse at an optimal time, in spite of the inherently stochastic expression of lysis proteins. These results illustrate the utility of the first-passage time framework for characterizing noise in the timing of intracellular events. Finally, analytical results and insights obtained here have broader implications for timing phenomenon in chemical kinetics, ecological modeling and statistical physics.

**FIG. 5.**
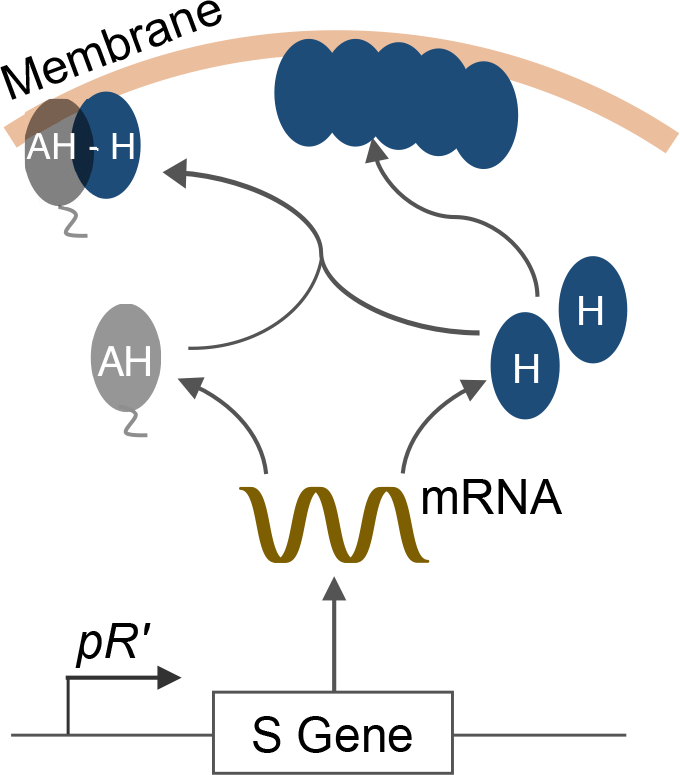
An incoherent feedforward circuit in *λ*’s lytic pathway. Bacteriophage *λ* lysis the infected host cell by expressing a membrane protein, holin (H). The protein slowly accumulates on the cell membrane over time and forms holes when a critical concentration threshold is reached. The mRNA encoding holin also expresses antiholin (AH), which binds to holin and prevents it from participating in hole formation creating a feedforward circuit.

## SUPPLEMENTARY INFORMATION

### S1. ON SOME PROPERTIES OF THE MATRIX A

In this section, we discuss some properties of the matrix **A** given in equation (9) of the main text.

#### S1-a. A is a Hurwitz matrix

The matrix **A** is given by

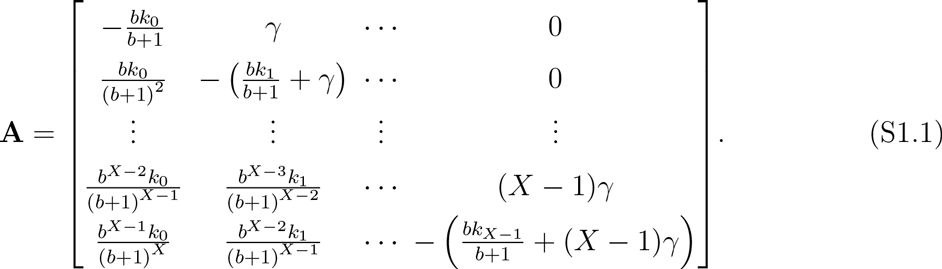

In order to prove that **A** is a Hurwitz matrix, we prove that the following two conditions hold true [84, pp. 48-49]:

1. The diagonal elements *a_ii_* < 0 for *i* = 1, 2,…, *X*,
2. 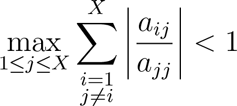.

Condition 1 here is satisfied here as 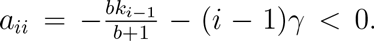. The left hand side of Condition 2 for any column *j* = 1, 2,…, *X* is

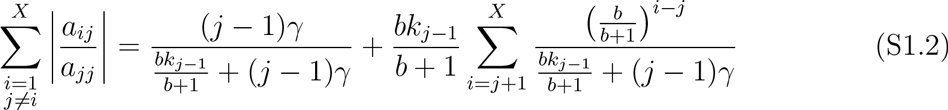

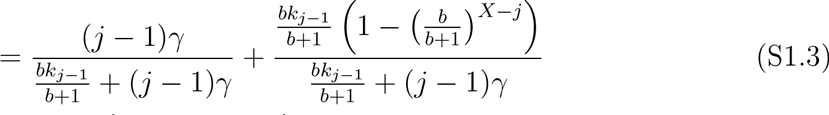

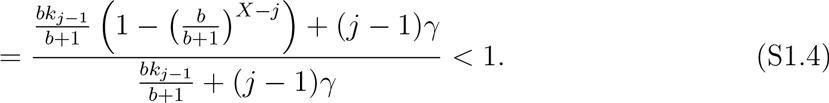

Thus, the matrix **A** is Hurwitz, i.e., the eigenvalues of **A** have negative real part.

#### S1-b. A is an invertible matrix

We will find the inverse of the matrix **A** thus proving that it is invertible.

Let us use **A**_0_ to denote the matrix **A** when *γ* = 0. The lower triangular matrix **A**_0_ is given by

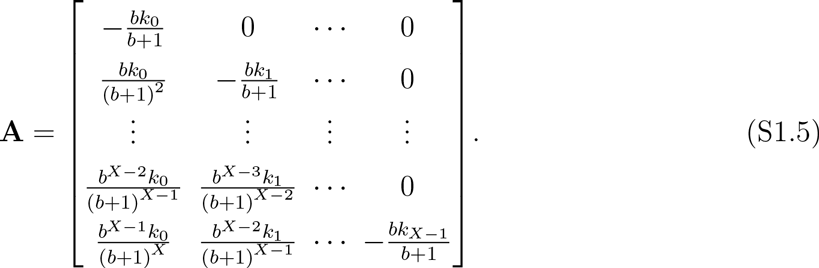

We claim that the inverse of **A**_0_ is given by the following matrix

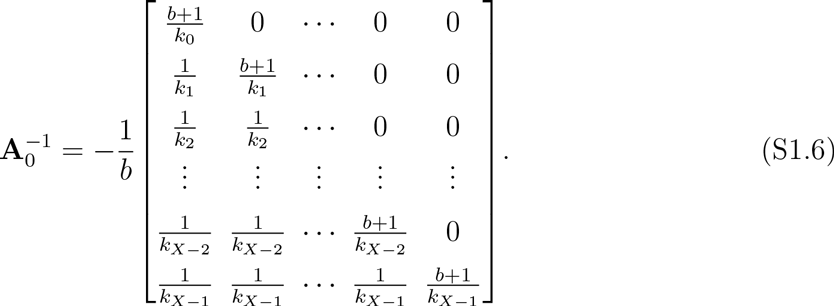

This claim can be quickly verified by multiplying the matrices which results in identity matrix. Next, to determine **A**^−1^, we observe that when *γ* ≠ 0, the matrix **A** can be written as

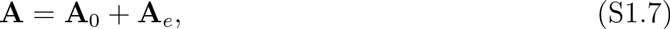

where **A**_e_ is given by

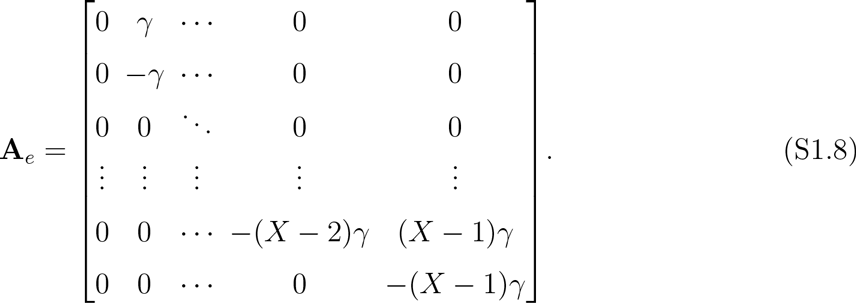

Therefore the inverse of the matrix **A** can be written as

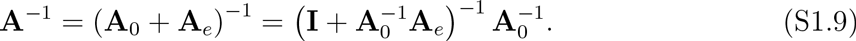

Note that **A**_e_ is a bidiagonal matrix with the *i^th^* diagonal element as − (*i* − 1)*γ*, 1 ≤ *i* ≤ *X* while the *j_th_* super-diagonal elements as *jγ*, 1 ≤ *j* ≤ *X* − 1. As we have already determined the expression of 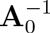 in (S1.6), the expression of 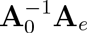 can be determined as

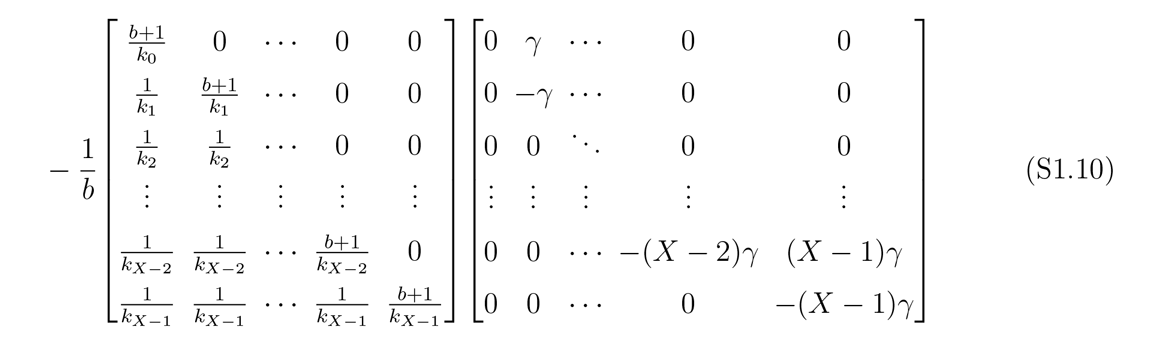

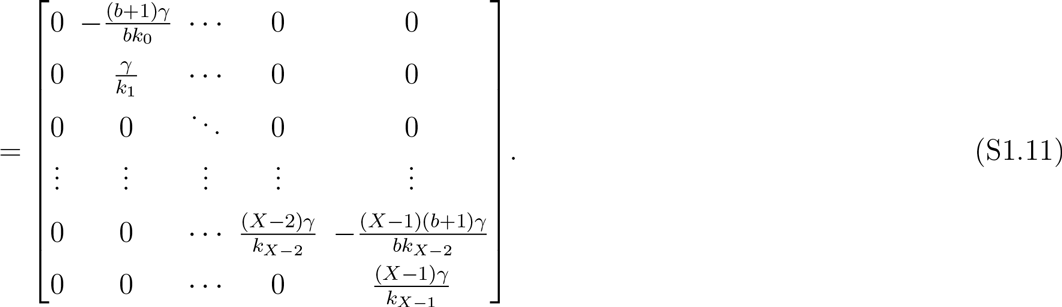

Thus, the matrix 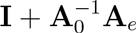 is a bidiagonal matrix with its diagonal elements 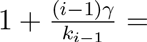 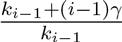 for *i* = 1, 2,…, *X*. The super diagonal elements are given by 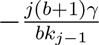 for *j* = 1,2,…, *X* − 1. Using the result for inverse of a bidiagonal matrix derived in [85], we can write the (*i*,*j*) element of inverse of 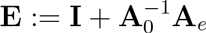 as follows

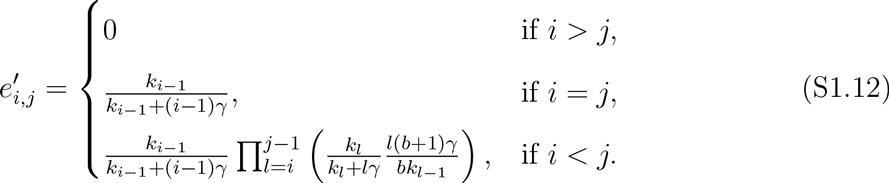

Alternatively, in matrix form

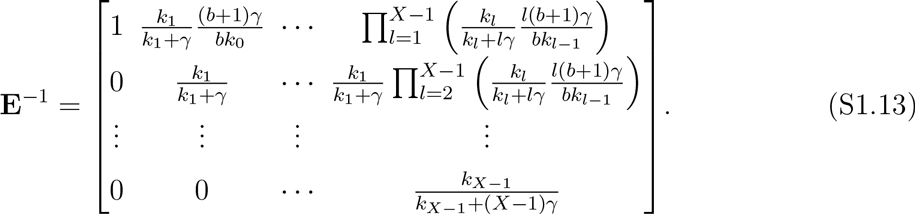

We can compute **A**^−1^ by calculating 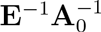. Here, we do not give explicit form of **A**^−1^ as it is not required for calculations in this document. Nevertheless, it validates that **A** is an invertible matrix.

### S2. EXPRESSION OF *m^th^* MOMENT OF FIRST-PASSAGE TIME

In this section, we make use of the properties discussed in the previous section to determine the moments of the first-passage time. As discussed in the main text (equation (12)), the distribution of first-passage time (*FPT*) is given by the following

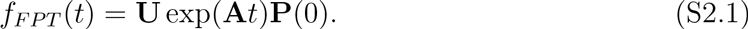

Therefore, a moment of *m^th^* order can be calculated as

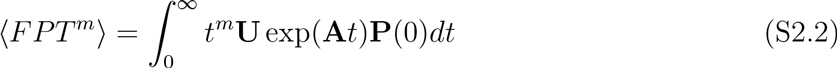

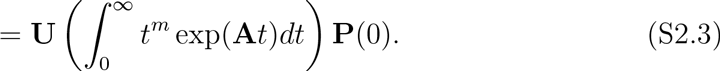

Let us consider 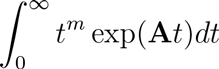. Integrating by parts

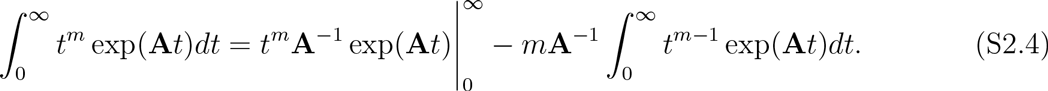

The first part of on the right hand side goes to zero as long as exp(**A***t*) goes to zero faster than the polynomial term *t^m^* which is true since the matrix **A** is a Hurwitz matrix, i.e., the eigen-values of **A** have negative real parts. Using *a_m_* as a notion to represent 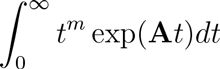 we can write the following recursive relationship

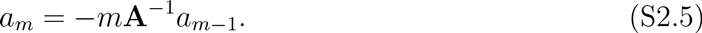

Thus, *a_m_* = (−1)^*m*^*m*!(**A**^−1^)^*m*^*a*_0_. Further,

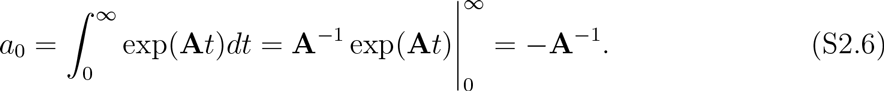

Therefore *a_m_* = (−1)^*m*+1^*m*!(**A**^−1^)^*m*+1^. Substituting this in (S2.3) gives the following for a general *m^th^* moment of FPT

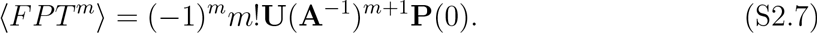

#### S2-a. Calculation of UA^−1^

As we saw in equation (S2.7), calculation of the moments will have a term of the form **UA**^−1^. Here we provide the calculation of this term.

Consider two matrices **G** and **H** such that **A**_e_ = **GH** where **G** is a *X* × *X* − 1 matrix

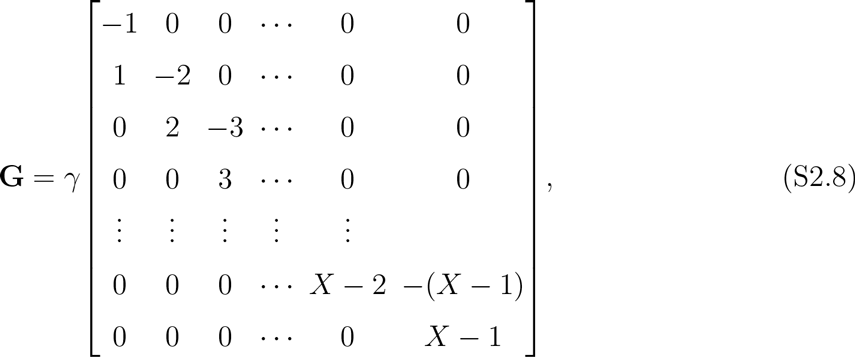

while **H** is a *X* − 1 × *X* matrix

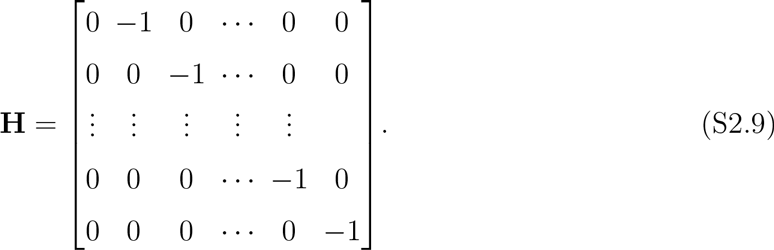

Using the matrix inversion lemma, **A**^−1^ can be written as

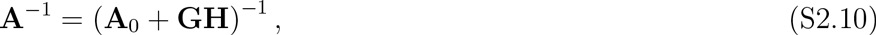

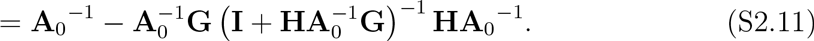

Let us look at the expression 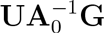.

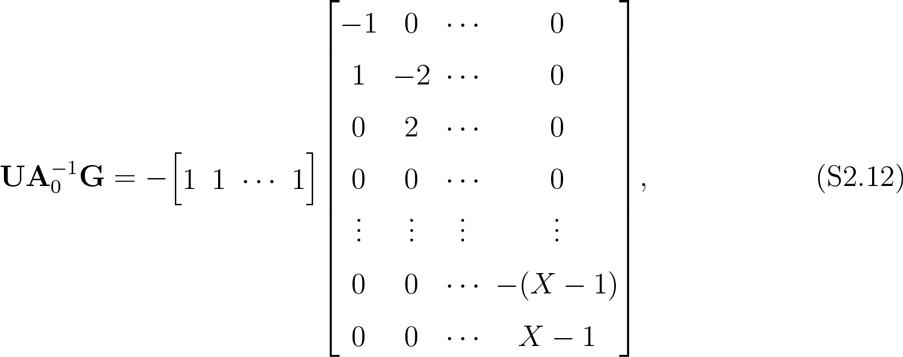

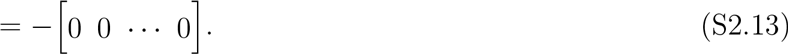

Therefore, we can conclude that **UA**^−1^ is in fact equal to 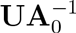 which could be calculated by multiplying **U** and 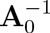.

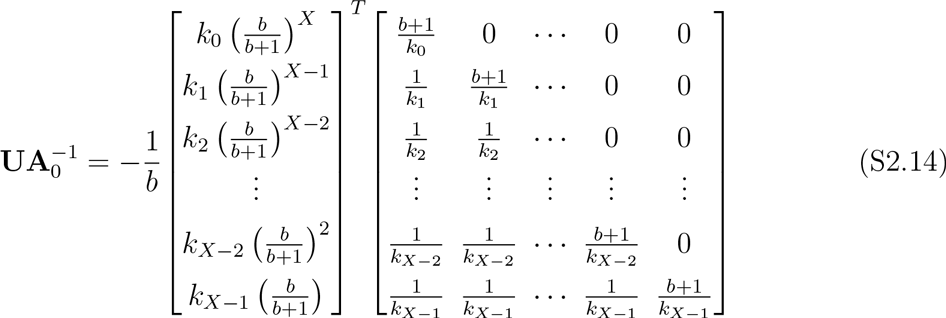

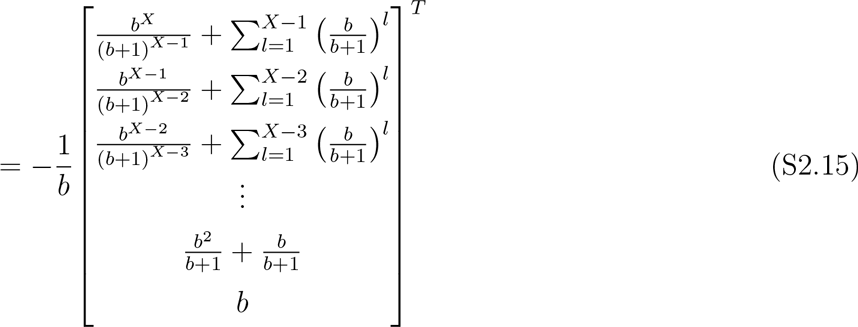

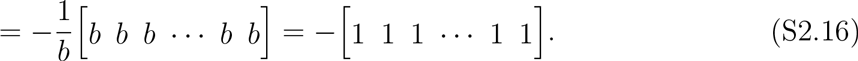

#### S2-b. Calculation of the first-two first-passage time moments

*Mean FPT* The mean FPT’s expression can be written as

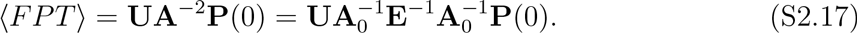

The expression of 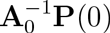 is just the first column of **A**_0_. Therefore

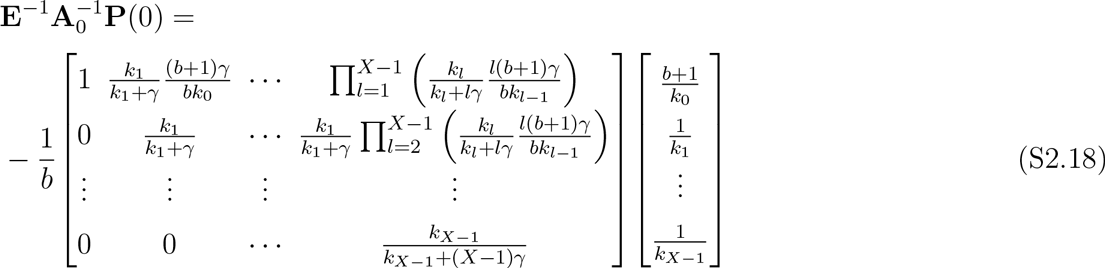

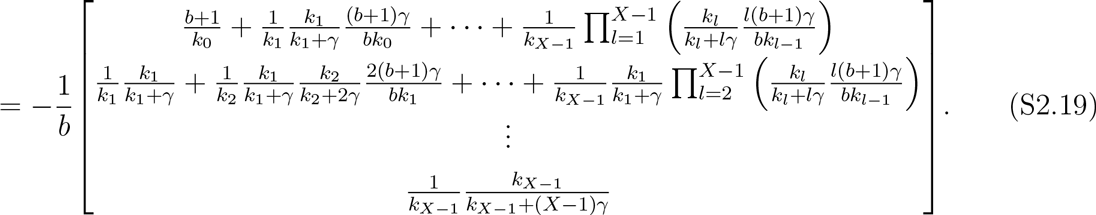

Since the vector 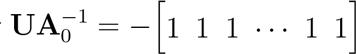, 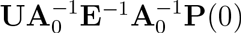 is essentially negative sum of the elements of 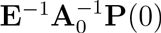. Therefore we have the expression of mean FPT is given by

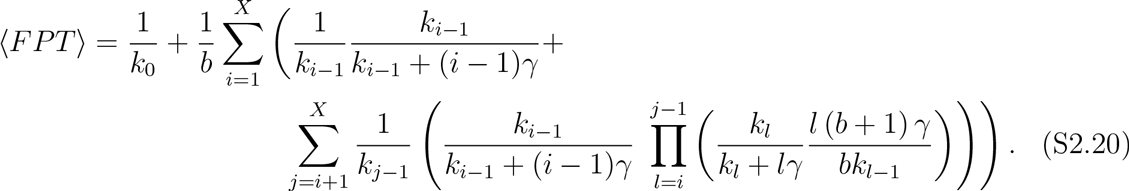

*a. Second order moment* The second order moment is given by

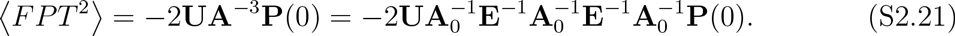

Let us use the notation *η_i_* defined as

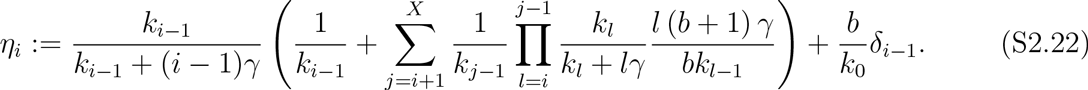

Using (S2.19), we can write

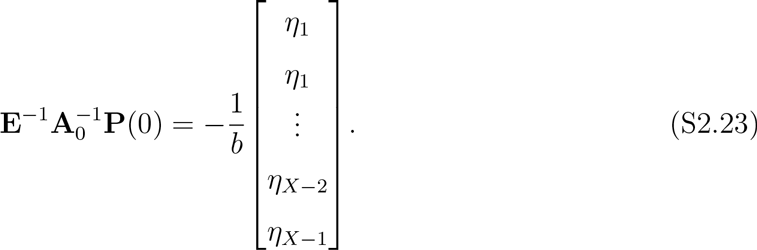

Therefore

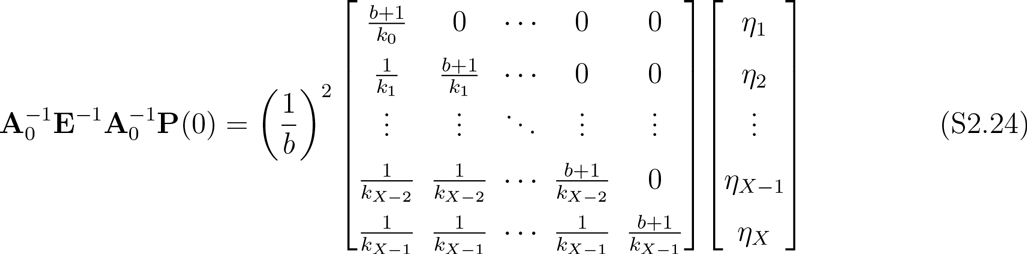

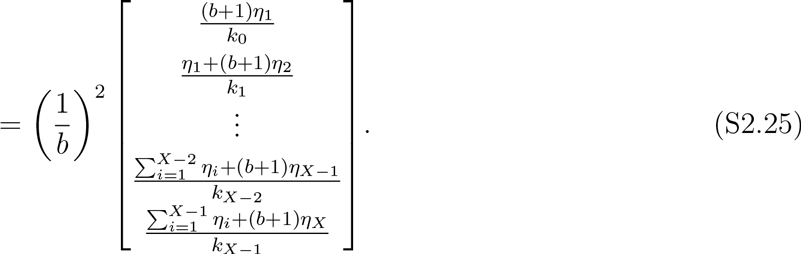

Using the notion 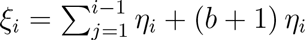, we can write 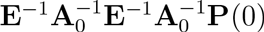 as

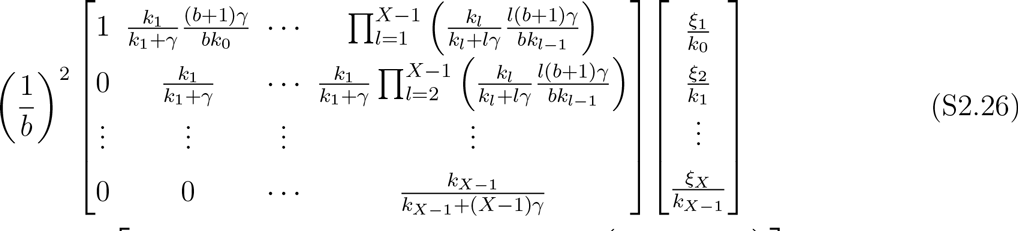

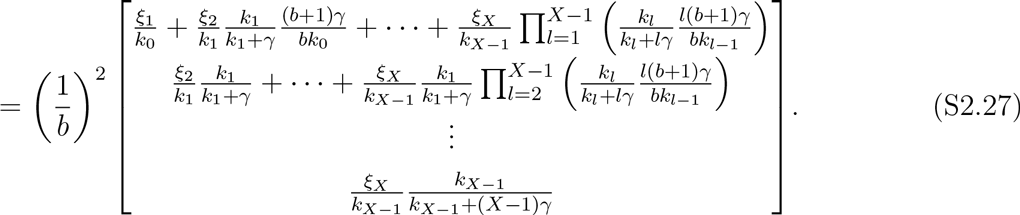

As 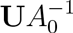 is merely negative summation of the elements of the column vector it pre-multiplies to, the second order moment of FPT can be given by following explicit formula

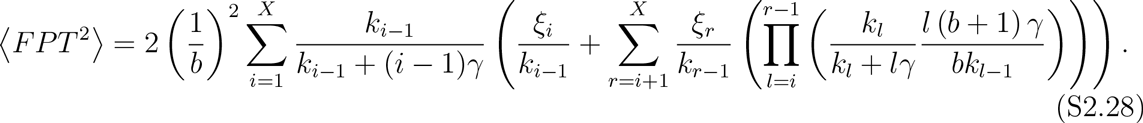

### S3. OPTIMAL FEEDBACK WHEN PROTEIN DOES NOT DEGRADE

As mentioned in the main text, our objective is to minimize ⟨*FPT*^2^⟩ such that ⟨*FPT*⟩ is fixed. For calculation purposes, we will denote this constraint as ⟨*FPT*⟩ = *t_opt_*. Let *m* represents the Lagrange’s multiplier, then we define the following objective function

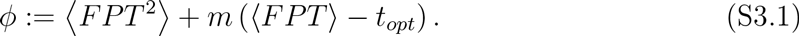

The optimization problem is solved in two steps. First, we determine the critical points. Second, we find the critical point corresponding to a global minimum.

Determining the critical points requires the following system of equations to be solved

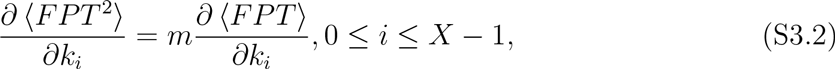

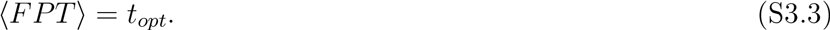

The expressions of the first-two moments of FPT when protein does not degrade are given by equation (16) in the main text. These are

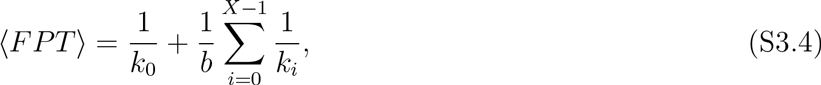

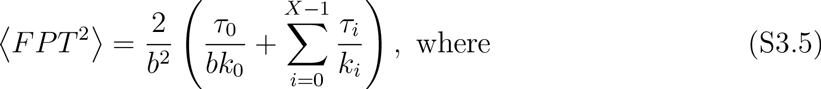

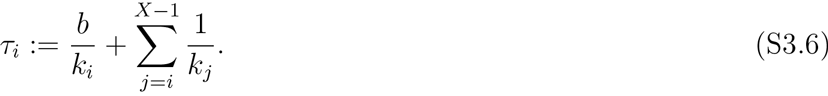

The optimization problem in equations (S3.2),(S3.3) requires calculation of the first order derivatives of ⟨*FPT*⟩ and ⟨*FPT*^2^⟩. The derivatives of ⟨*FPT*⟩ with respect to *k_i_*’s are given by

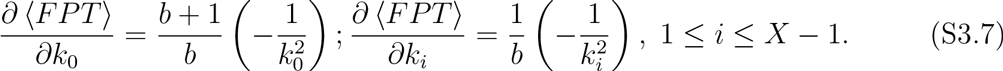

Similarly, the derivative of ⟨*FPT*^2^⟩ are

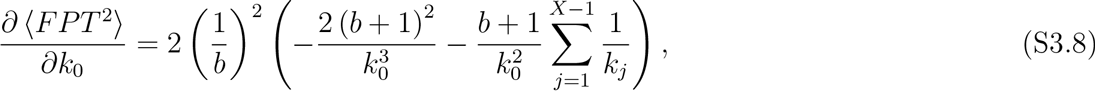

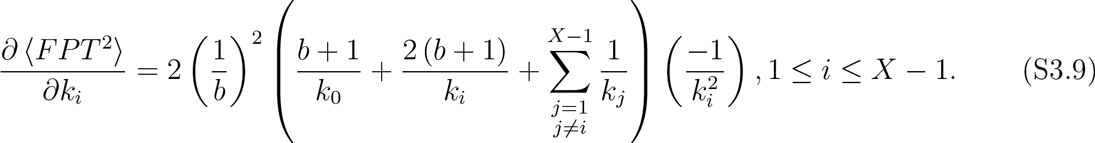

Therefore, the system of equations to be solved becomes

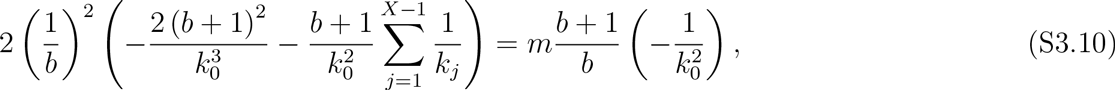

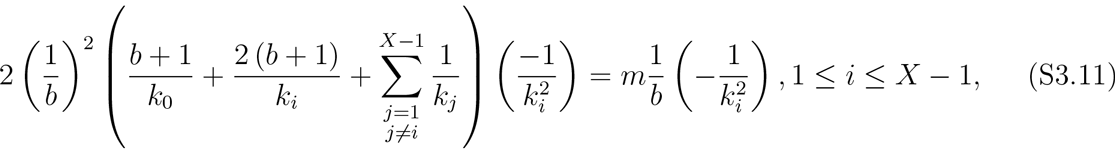

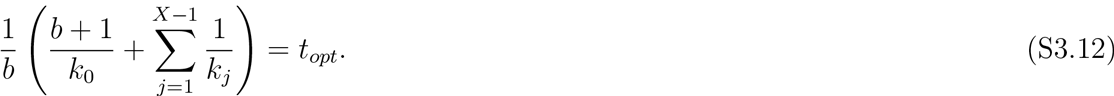

Furthermore, we want the solution such that none of the transcription rate is zero. This simplifies the system of equations to

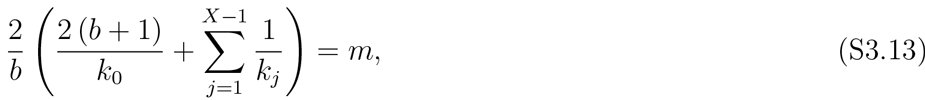

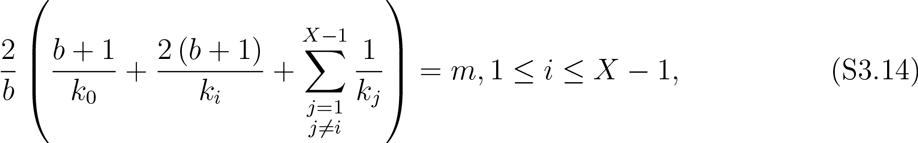

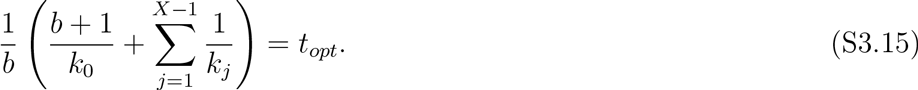

Solution to these equations is given by

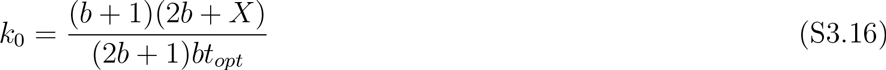

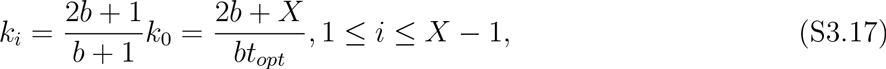

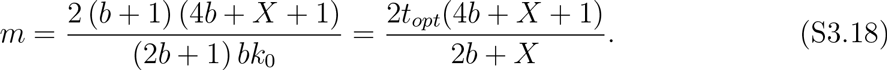

We have calculated the critical point for the optimization problem. However, it needs to be checked whether its an minimum or maximum. For this purpose, we consider the bordered Hessian as follows.

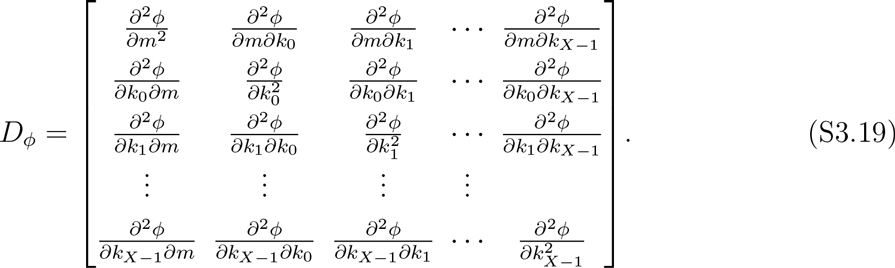

We will show that all the principal minors of this matrix are negative. To start with, let us first determine the second order derivatives of ⟨*FPT*⟩.

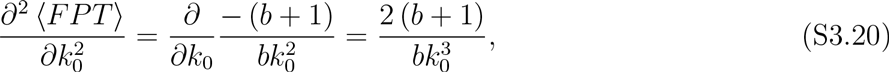

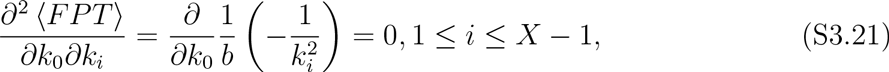

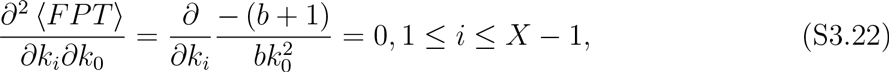

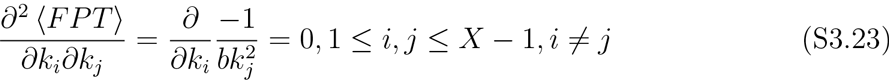

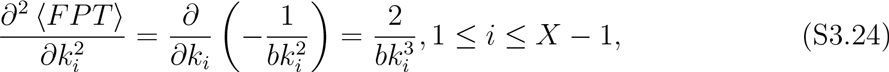

Similarly, the derivatives for ⟨*FPT*^2^⟩ are given by

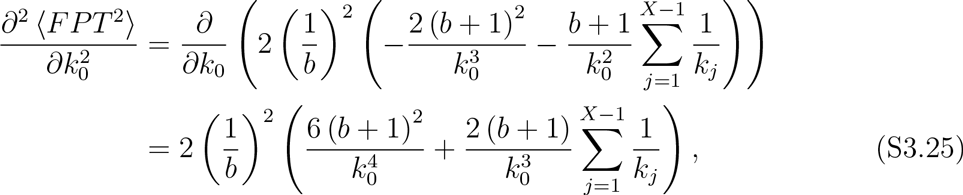

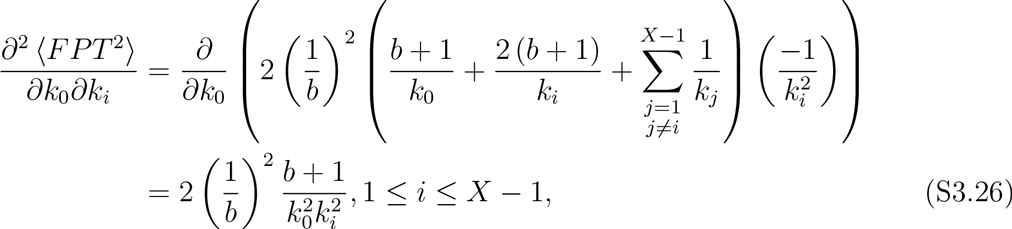

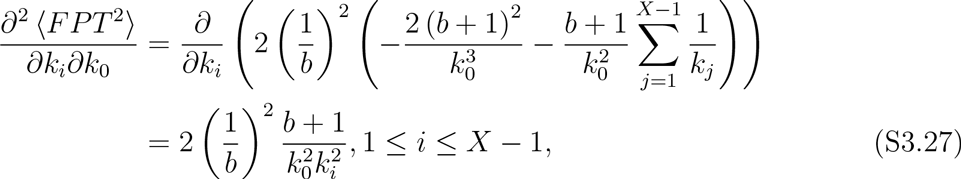

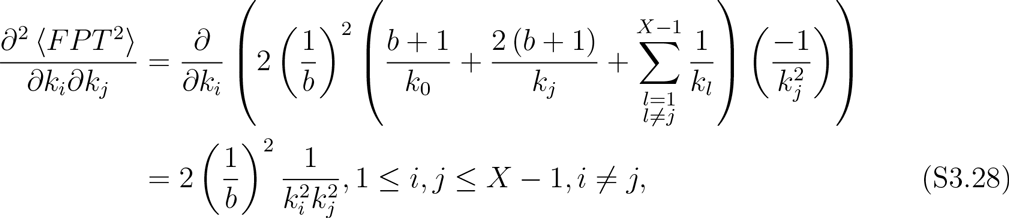

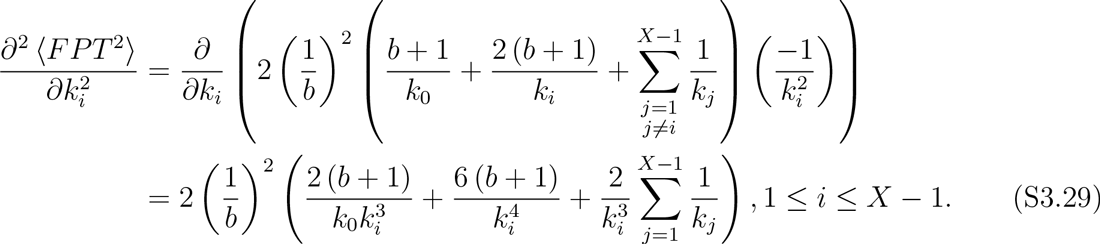

We can now determine the elements of the bordered Hessian matrix in equation (S3.19) computed at the solution given by equations (S3.16)–(S3.17).

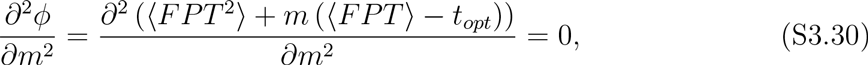

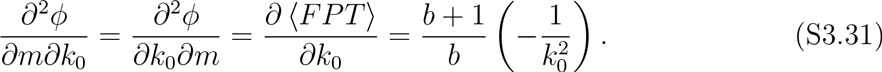

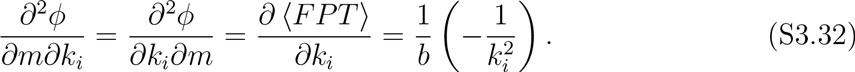

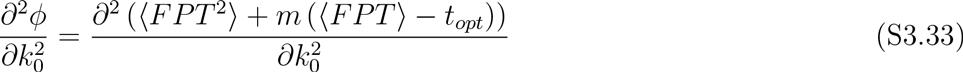

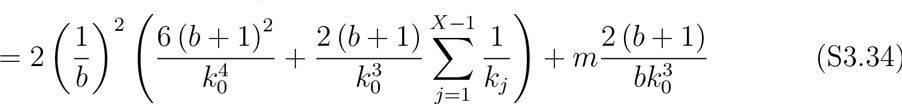

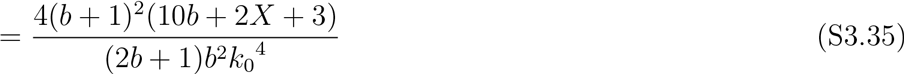

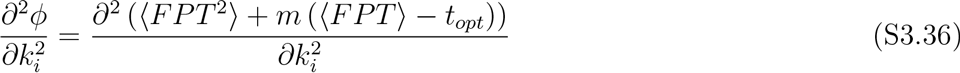

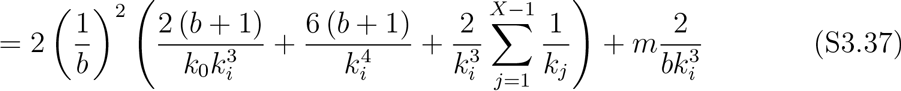

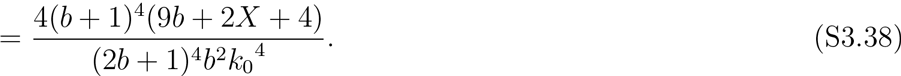

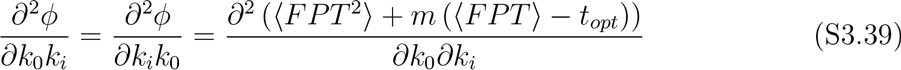

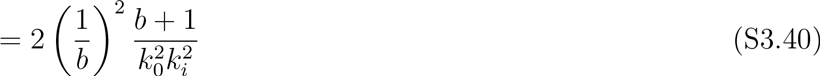

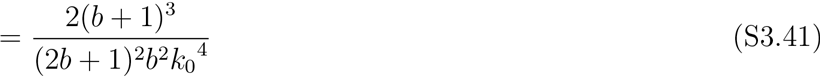

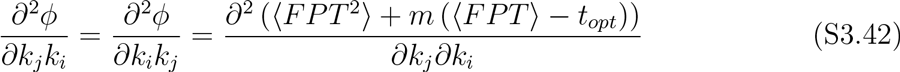

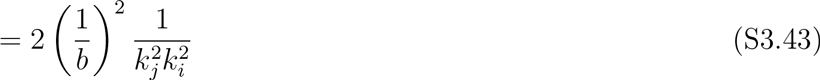

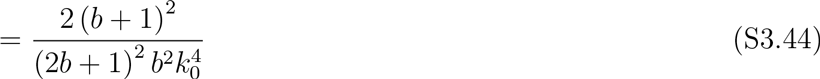

It can be noted elements of *D_ϕ_* are from a set six quantities. Defining

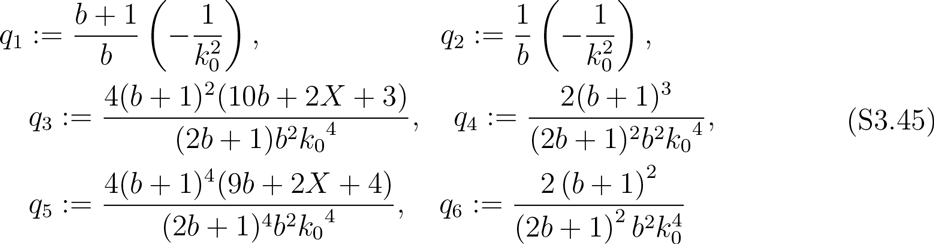

we can write *D_ϕ_* as

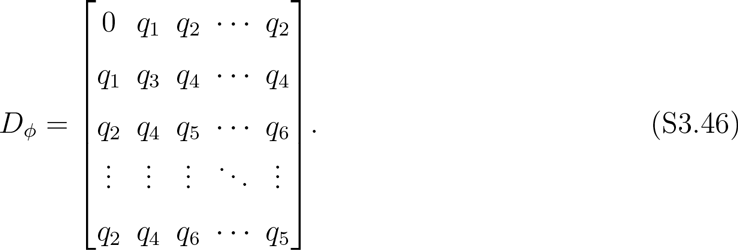

Let us denote by *K*(*n*) the principal minor of the matrix *D_ϕ_* of size *n* × *n*. It can be easily seen that*K*(1) = 0, 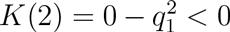 and 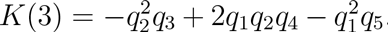. For 4 ≤ *n* ≤ *X*, we perform the following two elementary operations on *D_ϕ_*

- *col_r_* = *col_r_* − *col_r−_*
- *row_r_* = *row_i_*, − *row*_*r*−1_

for *r* = *n*,*n* − 1,…, 1. This yields

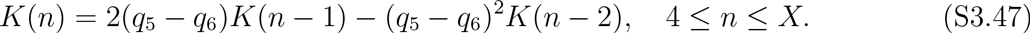

The solution to the above recursive equation is given by

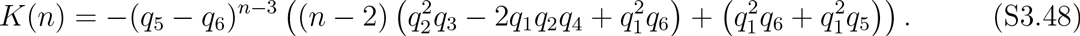

It can be easily checked that *K*(*n*) is negative because *q*_5_ > *q*_6_,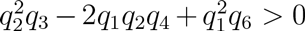 and 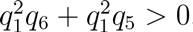. This proves that the critical point indeed corresponds to a minimum.

### S4. OPTIMAL FEEDBACK STRATEGY IN PRESENCE OF PROTEIN DEGRADATION

In previous sections, we have derived analytical expression of the optimal feedback strategy that minimizes ⟨*FPT*^2^⟩ such that ⟨*FPT*⟩ is constant. As the expressions of ⟨*FPT*⟩ and ⟨*FPT*^2^⟩ in equations (S2.20) and (S2.28) are too convoluted to solve for optimal transcription rates analytically, we take a numerical approach. For this purpose, we fixed the threshold *X* = 10 molecules, and mean burst size *b* = 1 molecules. Using numerical solvers, we searched the parameter space of the transcription rates *k_i_*,*i* ∈ {0,1,…, 9} for various values of the protein degradation rate *γ*. As shown in Fig. S4.1, when *γ* = 0, the transcription rates are equal except for the first one. This is consistent with the expressions in (S3.16)–(S3.17). Further, as the protein degradation rate is increased, the transcription rates first increase and then decrease, suggesting a mixed feedback strategy.

**FIG. S4.1.**
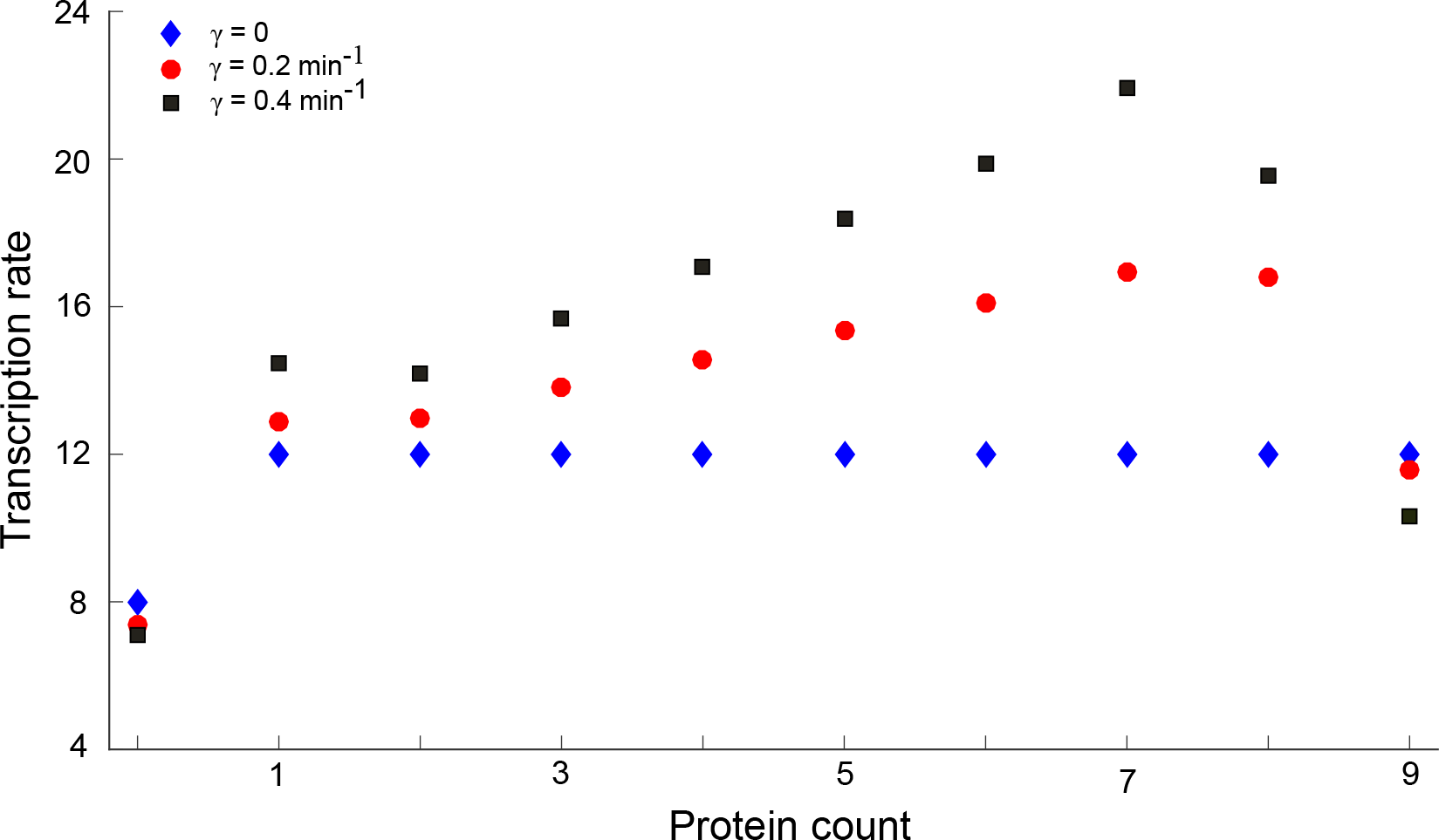
*Optimal feedback strategy for unstable protein*. The optimal transcription rates obtained via numerical optimization for different values of degradation rate are shown. The event threshold is assumed to be 10 molecules, and the mean *FPT* is constrained to be 1 minute.

To keep the results biologically meaningful, we assume that the feedbacks are implements as Hill functions. One simplest implementation of this would be a linear form of the feedback

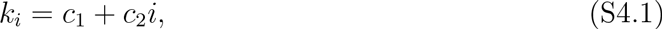

as considered in the analysis in the main text. Here, we show the results for having a nonlinear form of Hill functions wherein a positive feedback is implemented as below

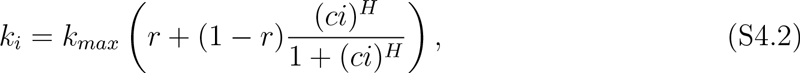

where *k_max_* is represents the maximum possible transcription rate, 0 < *r* < 1 is a constant corresponding to the minimum transcription rate (*k_max_r*), *c* is the feedback strength (note that *k_i_* = *k_max_*/2 when i=1/c), and *H* denotes the Hill coefficient. Similarly, a negative feedback is implemented via

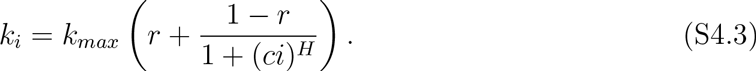

**FIG. S4.2.**
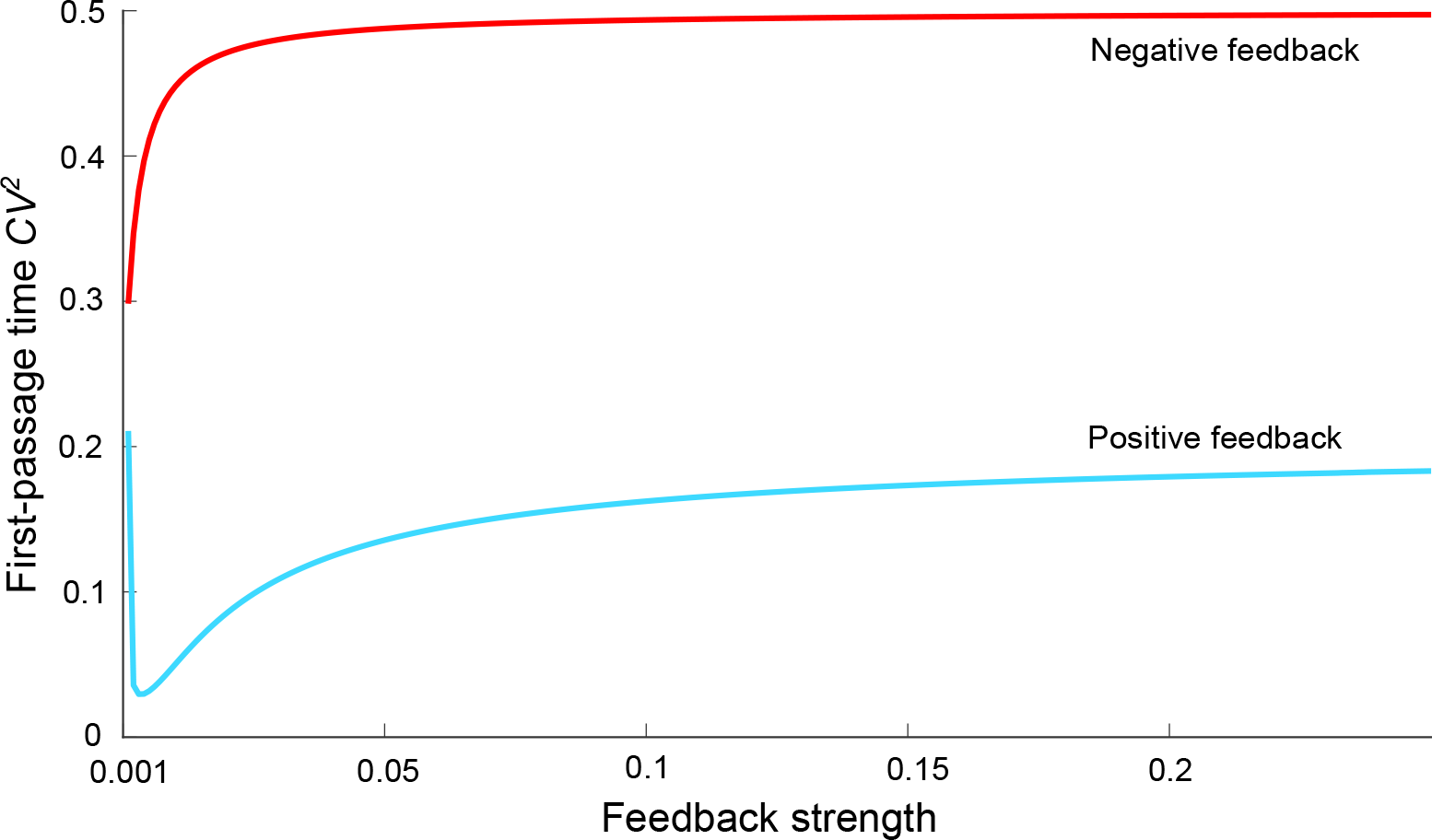
*Optimal feedback strategy in form of a Hill function*. The transcription rates are assumed to follow the Hill funciton forms as given by (S4.2) (positive feedback) and (S4.3) (negative feedback). For a negative feedback, 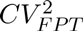 increases as the feedback strength increased. For a positive feedback, 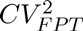 first decreases as feedback strength increases, hits a minimum at a certain feedback strength, and increases after that. The parameters used are *b* = 1 molecule, *r* = 0.05, and *H* = 1.

The results of these Hill function implementations are same as shown in the main text for the linear form of transcription rates. We show one example of this in Fig. S4.2. One can see that when the protein degradation is allowed, the negative feedback leads to increase in noise as its feedback strength is increased. For the positive feedback, the noise hits a minimum for a certain feedback strength. Here the mean *FPT* is kept constant by appropriately tuning the parameter *k_max_*.

One can see that there are more parameters in these forms of feedback, and various parameters (or their combinations) can be tuned to keep the mean fixed for a given feedback strength, the resulting analysis is complex. This is the rationale behind using the simplest possible form (linear) of the feedback which gives useful insight.

### S5. OPTIMAL FEEDBACK STRENGTH FOR LINEAR FORM OF FEEDBACK IN PRESENCE OF PROTEIN DEGRADATION

In this section, we explore the optimal feedback strength *c*_2_ as the protein degradation rate is varied. As in the main text, the feedbacks are assumed to follow a linear form given by *k_i_* = *c*_1_ ± *c*_2_*i*.

When the protein does not decay, as expected from theory, we get the optimal *c*_2_ = 0. As protein decay is considered, a positive feedback acts as the optimal feedback strategy. The optimal *c*_2_ multiplied by the mean burst size *b* takes values close to the degradation rate *γ* as shown in Fig. S5.1 (left). Interestingly, the protein trajectories generated by the optimal (positive) feedback in the case of protein degradation mimic those generated by the optimal feedback (no feedback) when protein did not decay. For their average dynamics to follow each other, a deterministic analysis reveals that as 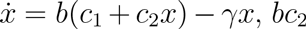, should be equal to *γ*. The slight difference between them appears to be due to the fact that a stochastic mean and a deterministic mean usually differ from each other.

**FIG. S5.1.**
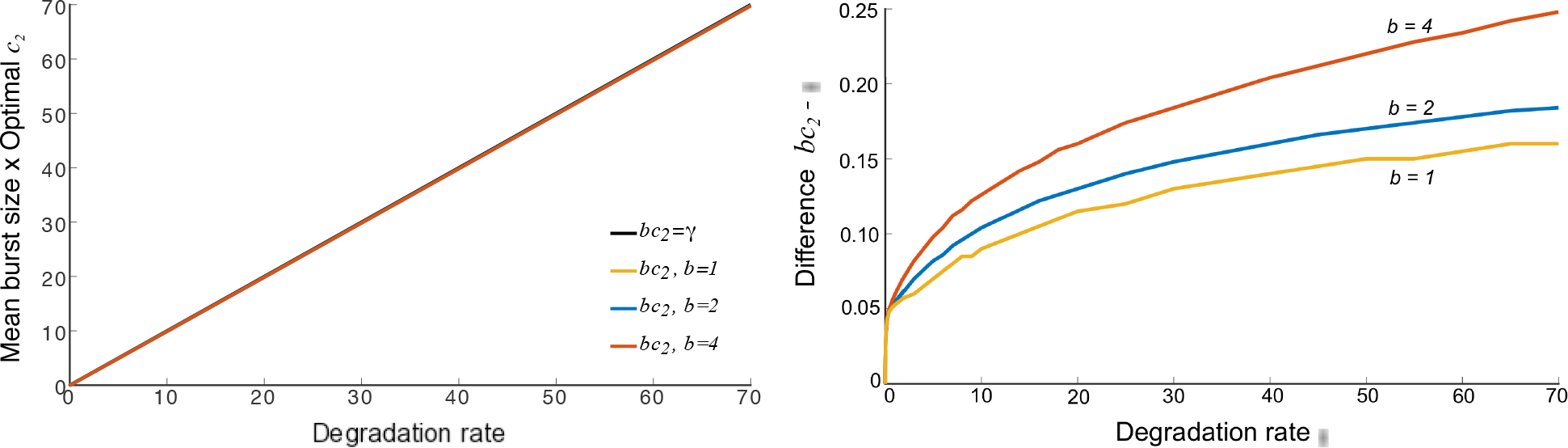
Difference between the degradation rate *γ* and optimal feedback strength *c_2_* multiplied by the mean burst size *b*. *Left*: The quantities are plotted for a range of degradation rates. It can be seen that they remain close to each other for increase in the degradation rate, and for several burst sizes. *Right*: The difference between the quantities is plotted as *γ* is changed. For a given mean burst size *b*, the difference *bc*_2_ − *γ* increases with increase in *γ*. For a given degradation rate, this difference also increases as the mean burst size *b* is increased.

We also explored the difference *bc*_2_ − *γ* as *γ* varies. Intriguingly, if the mean burst size *b* is kept constant, we observe that the difference increases for increase in *γ*. Further, for a given degradation rate, increasing *b* leads to increase in the difference *bc*_2_ − *γ*. The reason for this is not clear to us yet, though it appears that in this case the stochastic description of the dynamics shows more deviations from the deterministic dynamics.

### S6. OPTIMAL FEEDBACK FOR STABLE PROTEIN IN PRESENCE OF EXTRINSIC NOISE

In this section, assuming that the protein does not degrade, we investigate how the optimal regulation strategy deviates from a no feedback in presence of a static extrinsic noise. We consider two possibilities here: one, the extrinsic noise affects the mean burst size; two, the extrinsic noise affects the transcription rate. For the first case, we assume that the mean burst size is drawn from an arbitrary positive-valued distribution. The second case is analyzed by assuming that a factor *Z* multiplies with the transcription rates resulting in an effective transcription rate when *x*(*t*) = *i* to be *k_i_Z*.

#### S6-a. Optimal feedback when the mean burst size is drawn from a distribution

Let us assume the mean burst size a random variable with probability density function *f_b_*(*β*). Thus, the number of proteins in a burst are geometrically distributed with mean *b* where *b* itself is a random variable. The mean *FPT* can be computed as

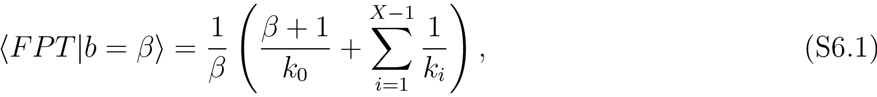

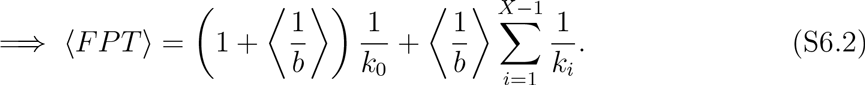

Similarly, the second order moment of *FPT* is given by

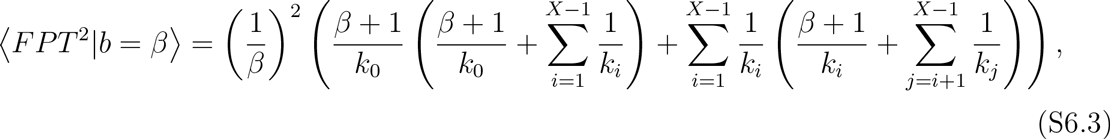

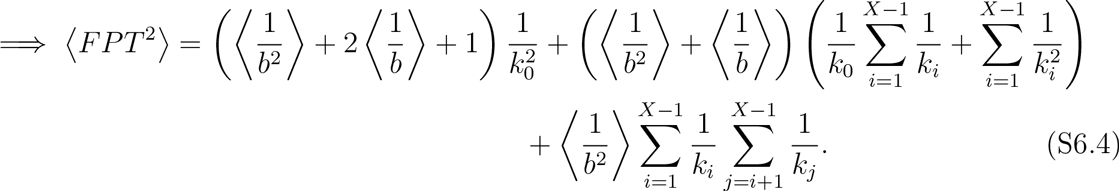

Defining 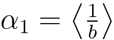 and 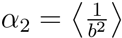, we have

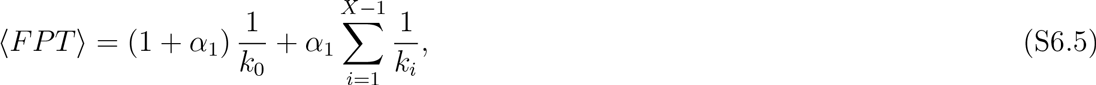

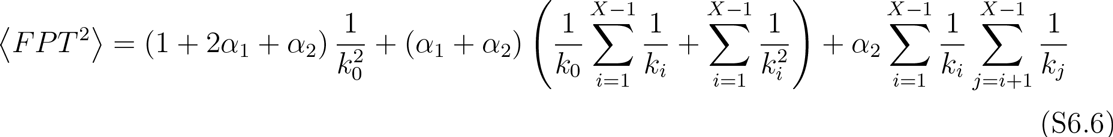

The derivatives of ⟨*FPT*⟩ with respect to *k_i_*’s are given by

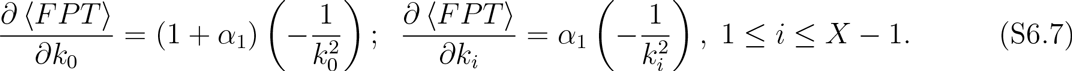

Similarly, the derivative of ⟨*FPT*^2^⟩ are

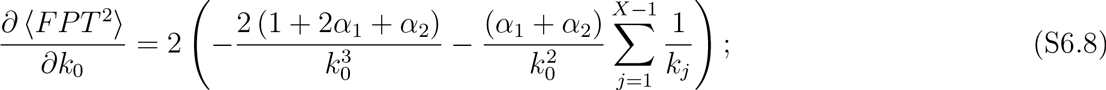

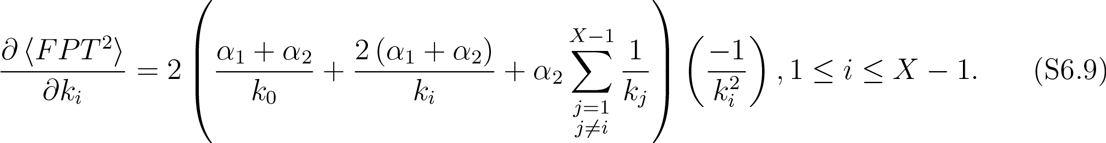

To find a critical point, we have to solve the following system of equations

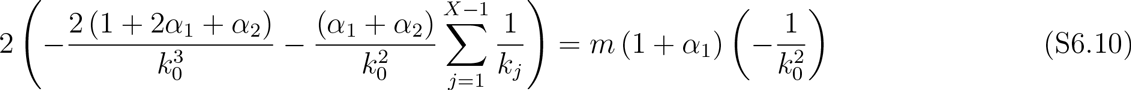

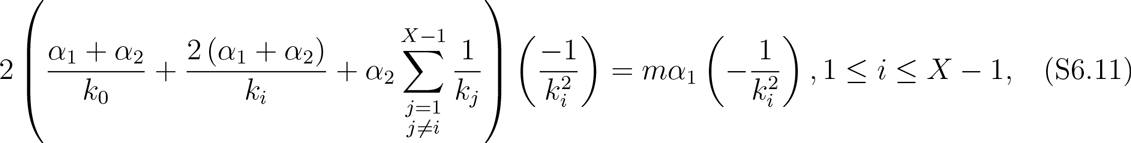

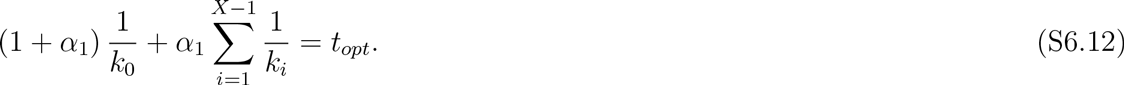

Assuming that *k*_0_, *k*_1_,… ≠ 0, we get

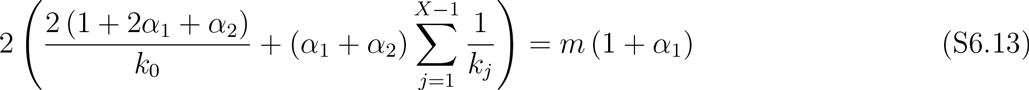

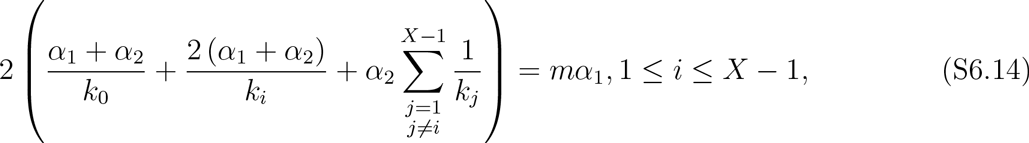

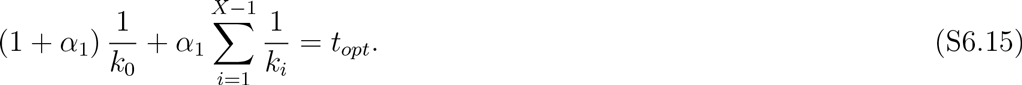

Solution to above system of equations gives

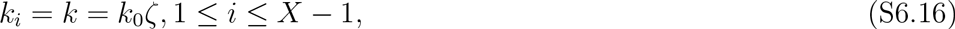

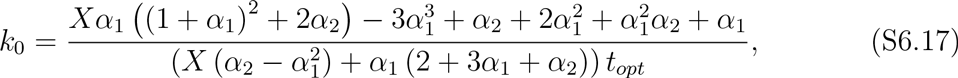

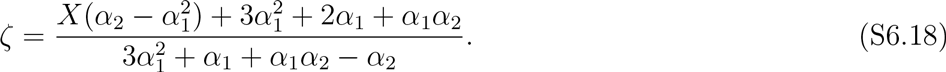

These equations reduce to our previous results of having a constant mean burst size *b* when *α*_1_ = 1/*b*, *α*_2_ = 1/*b*^2^ are used.

#### S6-b. Optimal regulation when extrinsic factor affects the transcription rate

We consider an extrinsic factor *Z* with a positive-valued arbitrary distribution *f_Z_*(*z*). This factor is assumed to be static, i.e., it does not vary over the time scale of the event of interest. Further we assume that it affects the transcription rates in a multiplicative fashion. The first-passage time mean in this case can be written as

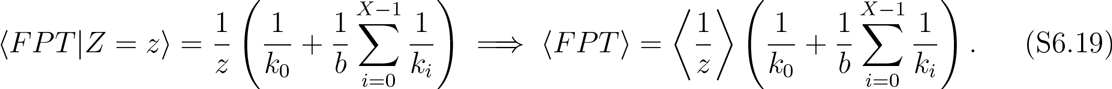

Likewise the second order moment can be written as

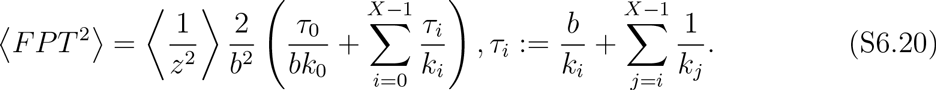

Solving the constrained optimization problem of minimizing ⟨*FPT*^2^⟩ constrained to ⟨*FPT*⟩ = *t_opt_* in this case simplifies to solving the following system of equations

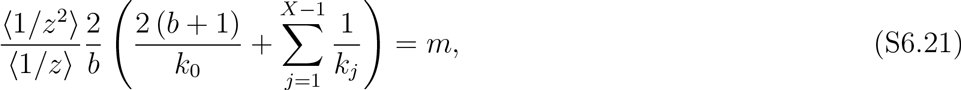

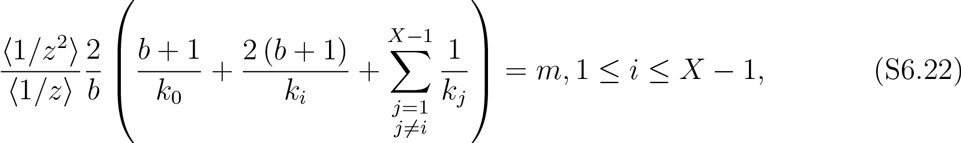

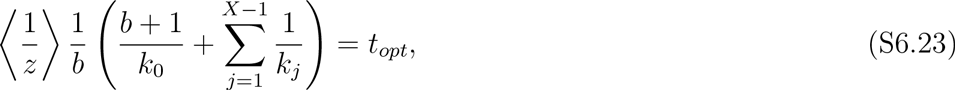

where *m* represents the Lagrange’s multiplier. Solution to these equations gives

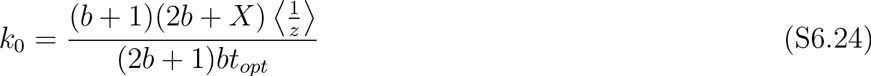

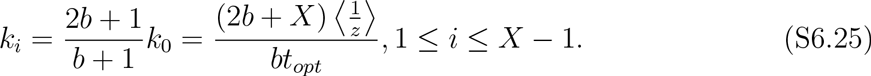

### S7. OPTIMAL FEEDBACK STRATEGY WHEN BURST SIZE IS DRAWN FROM A POISSON DISTRIBUTION

In the main paper, the mRNA degradation process is assumed to be memoryless (exponential), and consequently the burst of proteins is assumed to follow a geometric distribution. However, in the limit when the mRNA degradation process is deterministic, the burst size distribution becomes Poisson. For this reason, we investigate how the optimal feedback strategy changes in the case when the burst follows a Poisson distribution.

The production and degradation of the protein (similar to equation (2) in main text) is governed by following probabilities

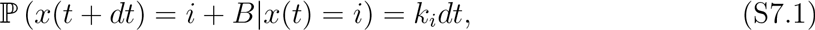

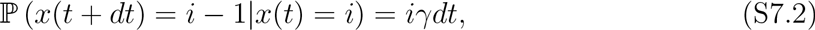

where the burst size *B* follows a Poisson distribution given by

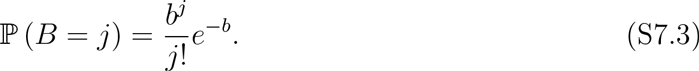

Here *b*, as before, represents the mean burst size, i.e., the average number of protein molecules produced by one mRNA.

**FIG. S7.1.**
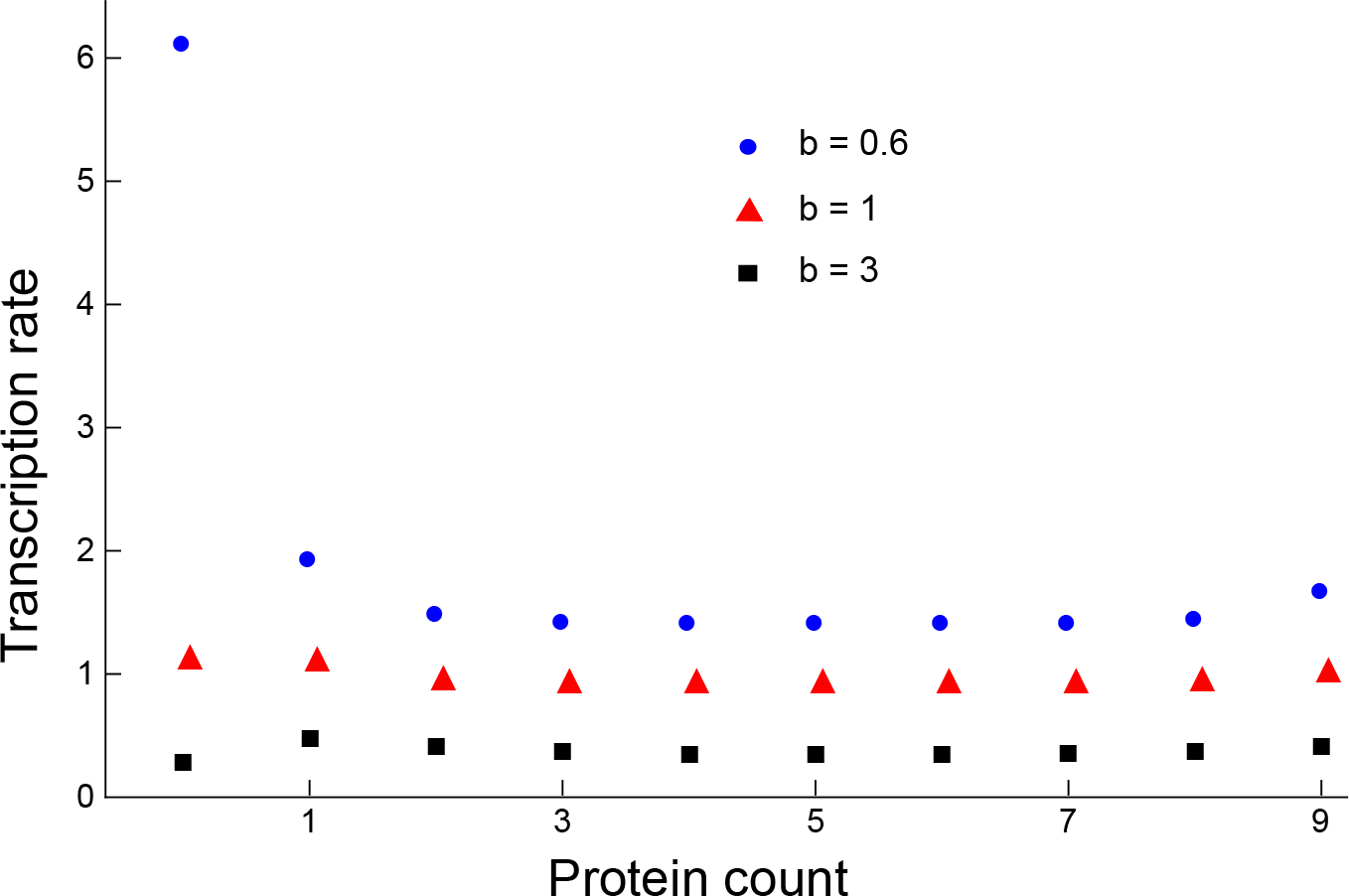
*Optimal feedback strategy for Poisson distributed burst size.* The optimal transcription rates obtained via numerical optimization for different values of mean burst size are shown. The event threshold is assumed to be 10 molecules, and the mean *FPT* is constrained to be 10 minutes.

One can carry out the first-passage time calculations in the same manner as done for the geometric burst size case. It turns out that the first two moments can be compactly written as

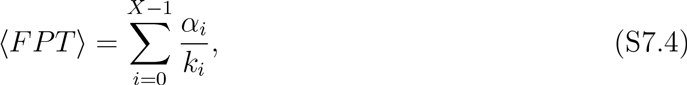

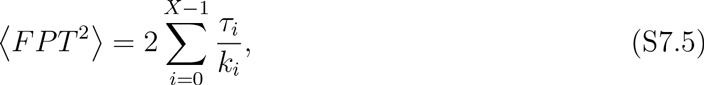

where

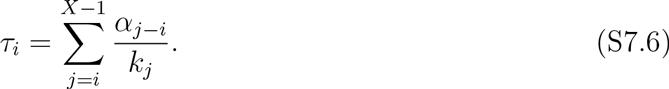

The coefficients *α_i_*, *i* ∈ {0,1, 2,… *X* − 1} are defined as

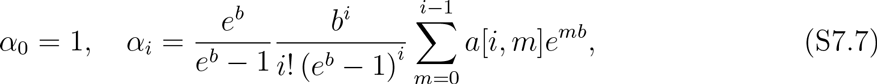

with *a*[*r*, *s*] represents an Eulerian number whose expression is

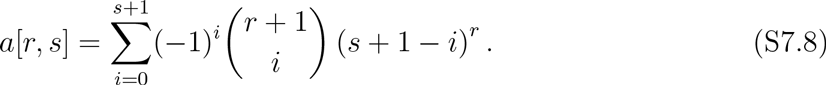

We performed numerical optimization with respect to parameters *k_i_*’s for threshold *X* = 10 to see the form of the optimal feedback strategy such that ⟨*FPT*^2^⟩ is minimized, with constraint ⟨*FPT*⟩ = *t_opt_* = 10 minutes. The results show that while the optimal transcription rates are not equal (i.e., no feedback mechanism in strict sense), they are fairly close to each other for mean burst sizes of 1 and 3 molecules. For mean burst size of 0.6 molecules, the first transcription rate when protein count is zero comes out to be significantly higher than other transcription rates which are more or less close to each other (Fig. S7.1). These results suggest that while the optimal feedback strategy deviates from a no feedback strategy with the underlying distribution of the burst size, it appears to remain close to a no feedback strategy.

## REFERENCES

[1] Katherine C Chen, Laurence Calzone, Attila Csikasz-Nagy, Frederick R Cross, Bela Novak, and John J Tyson. Integrative analysis of cell cycle control in budding yeast. Molecular biology of the cell, 15:3841–3862, 2004.

[2] James M Bean, Eric D Siggia, and Frederick R Cross. Coherence and timing of cell cycle start examined at single-cell resolution. Molecular Cell, 21(1):3–14, 2006.

[3] David L Satinover, David L Brautigan, and P Todd Stukenberg. Aurora-a kinase and inhibitor-2 regulate the cyclin threshold for mitotic entry in xenopus early embryonic cell cycles. Cell Cycle, 5:2268–2274, 2006.

[4] Xili Liu, Xin Wang, Xiaojing Yang, Sen Liu, Lingli Jiang, Yimiao Qu, Lufeng Hu, Qi Ouyang, and Chao Tang. Reliable cell cycle commitment in budding yeast is ensured by signal integration. eLife, 4:e03977, 2015.

[5] David T Champlin and James W Truman. Ecdysteroid control of cell proliferation during optic lobe neurogenesis in the moth manduca sexta. Development, 125:269–277, 1998.

[6] Takashi Koyama, Masafumi Iwami, and Sho Sakurai. Ecdysteroid control of cell cycle and cellular commitment in insect wing imaginal discs. Molecular and cellular endocrinology, 213:155–166, 2004.

[7] Karen Carniol, Patrick Eichenberger, and Richard Losick. A threshold mechanism governing activation of the developmental regulatory protein ctF in Bacillus subtilis. Journal of Biological Chemistry, 279:14860–14870, 2004.

[8] Patrick J Piggot and David W Hilbert. Sporulation of Bacillus subtilis. Current opinion in microbiology, 7:579–586, 2004.

[9] Sabrina L Spencer, Suzanne Gaudet, John G Albeck, John M Burke, and Peter K Sorger. Non-genetic origins of cell-to-cell variability in TRAIL-induced apoptosis. Nature, 459(7245):428–432, 2009.

[10] Jérémie Roux, Marc Hafner, Samuel Bandara, Joshua J Sims, Hannah Hudson, Diana Chai, and Peter K Sorger. Fractional killing arises from cell-to-cell variability in overcoming a caspase activity threshold. Molecular systems biology, 11:803, 2015.

[11] M Kracikova, G Akiri, A George, R Sachidanandam, and SA Aaronson. A threshold mechanism mediates p53 cell fate decision between growth arrest and apoptosis. Cell Death & Differentiation, 20:576–588, 2013.

[12] Helen K Salz. Male or female? the answer depends on when you ask. PLoS biology, 5, 2007.

[13] Yifat Goldschmidt, Evgeny Yurkovsky, Amit Reif, Roni Rosner, Amit Akiva, and Iftach Nachman. Control of relative timing and stoichiometry by a master regulator. PLOS ONE, 10:1–14, 2015.

[14] Harley H McAdams and Adam Arkin. Stochastic mechanisms in gene expression. Proceedings of the National Academy of Sciences, 94(3):814–819, 1997.

[15] Iftach Nachman, Aviv Regev, and Sharad Ramanathan. Dissecting timing variability in yeast meiosis. Cell, 131(3):544–556, 2007.

[16] Juan M Pedraza and Johan Paulsson. Random timing in signaling cascades. Molecular Systems Biology, 3(1), 2007.

[17] Long Cai and Nir Friedmanand X. Sunney Xie. Stochastic protein expression in individual cells at the single molecule level. Nature, 440:358–362, September 2006.

[18] Daniel K. Wells, William L. Kath, and Adilson E. Motter. Control of stochastic and induced switching in biophysical networks. Physical Review X, 5:031036, 2015.

[19] Jonathan M Raser and Erin K O’Shea. Noise in gene expression: origins, consequences, and control. Science, 309(5743):2010–2013, 2005.

[20] Arjun Raj and Alexander van Oudenaarden. Nature, nurture, or chance: stochastic gene expression and its consequences. Cell, 135(2):216–226, 2008.

[21] Mads Ksrn, Timothy C Elston, William J Blake, and James J Collins. Stochasticity in gene expression: from theories to phenotypes. Nature Reviews Genetics, 6(6):451–464, 2005.

[22] Adam M Corrigan, Edward Tunnacliffe, Danielle Cannon, and Jonathan R Chubb. A continuum model of transcriptional bursting. eLife, 5:e13051, 2016.

[23] R. D. Dar, B. S. Razooky, A. Singh, T. V. Trimeloni, J. M. McCollum, C. D. Cox, M. L. Simpson, and L. S. Weinberger. Transcriptional burst frequency and burst size are equally modulated across the human genome. Proceedings of the National Academy of Sciences, 109:17454–17459, 2012.

[24] D. M. Suter, N. Molina, D. Gatfield, K. Schneider, U. Schibler, and F. Naef. Mammalian genes are transcribed with widely different bursting kinetics. Science, 332:472–474, 2011.

[25] Michael Hinczewski and D. Thirumalai. Cellular signaling networks function as generalized wiener-kolmogorov filters to suppress noise. Physical Review X, 4:041017, 2014.

[26] A. Singh, Brandon Razooky, Chris D. Cox, Michael L. Simpson, and Leor S. Weinberger. Transcriptional bursting from the HIV-1 promoter is a significant source of stochastic noise in HIV-1 gene expression. Biophysical Journal, 98:L32–L34, 2010.

[27] Amnon Amir, Oren Kobiler, Assaf Rokney, Amos B Oppenheim, and Joel Stavans. Noise in timing and precision of gene activities in a genetic cascade. Molecular Systems Biology, 3(1), 2007.

[28] John J Dennehy and Nang Wang. Factors influencing lysis time stochasticity in bacteriophage A. BMC Microbiology, 11(1):174, 2011.

[29] Evgeny Yurkovsky and Iftach Nachman. Event timing at the single-cell level. Briefings in Functional Genomics, 12(2):90–98, 2013.

[30] DAJ Middleton, AR Veitch, and RM Nisbet. The effect of an upper limit to population size on persistence time. Theoretical Population Biology, 48:277–305, 1995.

[31] Johan Grasman and Reinier HilleRisLambers. On local extinction in a metapopulation. Ecological Modelling, 103:71–80, 1997.

[32] TJ Newman, Jean-Baptiste Ferdy, and C Quince. Extinction times and moment closure in the stochastic logistic process. Theoretical Population Biology, 65:115–126, 2004.

[33] Per Fauchald and Torkild Tveraa. Using first-passage time in the analysis of area-restricted search and habitat selection. Ecology, 84:282–288, 2003.

[34] Otso Ovaskainen and Baruch Meerson. Stochastic models of population extinction. Trends in Ecology & Evolution, 25:643–652, 2010.

[35] George H Weiss and Menachem Dishon. On the asymptotic behavior of the stochastic and deterministic models of an epidemic. Mathematical Biosciences, 11:261–265, 1971.

[36] Alun L Lloyd and Robert M May. How viruses spread among computers and people. Science, 292:1316–1317, 2001.

[37] D Volovik and S Redner. First-passage properties of bursty random walks. Journal of Statistical Mechanics: Theory and Experiment, 2010:P06018, 2010.

[38] Daniel A Charlebois, Nezar Abdennur, and Mads Kaern. Gene expression noise facilitates adaptation and drug resistance independently of mutation. Physical Review Letters, 107:218101, 2011.

[39] Tom Chou and Yu Wang. Fixation times in differentiation and evolution in the presence of bottlenecks, deserts, and oases. Journal of Theoretical Biology, 372:65–73, 2015.

[40] Golan Bel, Brian Munsky, and Ilya Nemenman. The simplicity of completion time distributions for common complex biochemical processes. Physical biology, 7:016003, 2010.

[41] Srividya Iyer-Biswas and Anton Zilman. First-Passage Processes in Cellular Biology, pages 261–306. John Wiley & Sons Inc, 2016.

[42] Wei Dai, Anirvan M. Sengupta, and Ronald M. Levy. First passage times, lifetimes, and relaxation times of unfolded proteins. Physical Review Letters, 115:048101, 2015.

[43] Khem Raj Ghusinga and Abhyudai Singh. First-passage time calculations for a gene expression model. IEEE Conference on Decision and Control, pages 3047–3052, 2014.

[44] Attila Becskei and Luis Serrano. Engineering stability in gene networks by autoregulation. Nature, 405(6786):590–593, 2000.

[45] Uri Alon. Network motifs: theory and experimental approaches. Nature Reviews Genetics, 8(6):450–461, 2007.

[46] Abhyudai Singh and Joao Pedro Hespanha. Evolution of gene auto-regulation in the presence of noise. Systems Biology, IET, 3(5):368–378, 2009.

[47] Abhyudai Singh. Negative feedback through mrna provides the best control of gene-expression noise. IEEE Transactions on NanoBioscience, 10(3):194–200, 2011.

[48] Ioannis Lestas, Glenn Vinnicombe, and Johan Paulsson. Fundamental limits on the suppression of molecular fluctuations. Nature, 467:174–178, 2010.

[49] Yi Tao, Xiudeng Zheng, and Yuehua Sun. Effect of feedback regulation on stochastic gene expression. Journal of Theoretical Biology, 247(4):827–836, 2007.

[50] Abhyudai Singh and Joao P Hespanha. Optimal feedback strength for noise suppression in autoregulatory gene networks. Biophysical Journal, 96(10):4013–4023, 2009.

[51] T. B. Kepler and T. C. Elston. Stochasticity in transcriptional regulation: Origins, consequences, and mathematical representations. Biophysical Journal, 81:3116–3136, 2001.

[52] Sara Hooshangi and Ron Weiss. The effect of negative feedback on noise propagation in transcriptional gene networks. CHAOS, 16, 2006.

[53] Margaritis Voliotis and Clive G. Bowsher. The magnitude and colour of noise in genetic negative feedback systems. Nucleic Acids Research, 2012.

[54] Abhyudai Singh and Mohammad Soltani. Quantifying intrinsic and extrinsic variability in stochastic gene expression models. PloS One, 8(12):e84301, 2013.

[55] Vahid Shahrezaei, Julien F. Ollivier, and Peter S. Swain. Colored extrinsic fluctuations and stochastic gene expression. Molecular Systems Biology, 4, 2008.

[56] Peter S Swain, Michael B Elowitz, and Eric D Siggia. Intrinsic and extrinsic contributions to stochasticity in gene expression. Proceedings of the National Academy of Sciences, 99(20):12795–12800, 2002.

[57] Andreas Hilfinger and Johan Paulsson. Separating intrinsic from extrinsic fluctuations in dynamic biological systems. Proceedings of the National Academy of Sciences, 108:12167–12172, 2011.

[58] David R. Rigney. Stochastic model of constitutive protein levels in growing and dividing bacterial cells. Journal of Theoretical Biology, 76(4):453 – 480, 1979.

[59] Johan Paulsson. Models of stochastic gene expression. Physics of Life Reviews, 2(2):157–175, 2005.

[60] Nir Friedman, Long Cai, and X Sunney Xie. Linking stochastic dynamics to population distribution: an analytical framework of gene expression. Physical Review Letters, 97(16):168302, 2006.

[61] Vlad Elgart, Tao Jia, Andrew T Fenley, and Rahul Kulkarni. Connecting protein and mrna burst distributions for stochastic models of gene expression. Physical biology, 8(4):046001, 2011.

[62] Niraj Kumar, Abhyudai Singh, and Rahul V. Kulkarni. Transcriptional bursting in gene expression: Analytical results for genera stochastic models. PLOS Computational Biology, 11:e1004292, 2015.

[63] Otto G Berg. A model for the statistical fluctuations of protein numbers in a microbial population. Journal of Theoretical Biology, 71(4):587–603, 1978.

[64] Ji Yu, Jie Xiao, Xiaojia Ren, Kaiqin Lao, and X Sunney Xie. Probing gene expression in live cells, one protein molecule at a time. Science, 311(5767):1600–1603, 2006.

[65] Vahid Shahrezaei and Peter S Swain. Analytical distributions for stochastic gene expression. Proceedings of the National Academy of Sciences, 105(45):17256–17261, 2008.

[66] Donald A McQuarrie. Journal of applied probability, 4:413–478, 1967.

[67] NG Van Kampen. Stochastic Processes in Physics and Chemistry. Elsevier, 2011.

[68] DT Gillespie. Exact stochastic simulation of coupled chemical reactions. J Phys Chem, 81:2340–2361, 1977.

[69] Abhyudai Singh. Transient changes in intercellular protein variability identify sources of noise in gene expression. Biophysical Journal, 107:2214–2220, 2014.

[70] Rebecca White, Shinobu Chiba, Ting Pang, Jill S Dewey, Christos G Savva, Andreas Holzenburg, Kit Pogliano, and Ry Young. Holin triggering in real time. Proceedings of the National Academy of Sciences, 108(2):798–803, 2011.

[71] Ryland Young. Phage lysis: Three steps, three choices, one outcome. Journal of Microbiology, 52:243–258, 2014.

[72] Ing-Nang Wang, David L Smith, and Ry Young. Holins: the protein clocks of bacteriophage infections. Annual Reviews in Microbiology, 54(1):799–825, 2000.

[73] Ing-Nang Wang, Daniel E Dykhuizen, and Lawrence B Slobodkin. The evolution of phage lysis timing. Evolutionary Ecology, 10(5):545–558, 1996.

[74] Ing-Nang Wang. Lysis timing and bacteriophage fitness. Genetics, 172(1):17–26, 2006.

[75] Richard H Heineman and James J Bull. Testing optimality with experimental evolution: lysis time in a bacteriophage. Evolution, 61(7):1695–1709, 2007.

[76] Yongping Shao and Ing-Nang Wang. Bacteriophage adsorption rate and optimal lysis time. Genetics, 180(1):471–482, 2008.

[77] Juan A Bonachela and Simon A Levin. Evolutionary comparison between viral lysis rate and latent period. Journal of Theoretical Biology, 345:32–42, 2014.

[78] Chung-Yu Chang, Kiebang Nam, and RY Young. S gene expression and the timing of lysis by bacteriophage lambda. Journal of bacteriology, 177(11):3283–3294, 1995.

[79] Yongping Shao and Ing-Nang Wang. Effect of late promoter activity on bacteriophage A fitness. Genetics, 181(4):1467–1475, 2009.

[80] Angelika Grundling, David L. Smith, Udo Blasi, and Ry Young. Dimerization between the holin and holin inhibitor of phage A. Journal of Bacteriology, 182:6075–6081, 2000.

[81] Udo Blasi and Ry Young. Two beginnings for a single purpose: the dual-start holins in the regulation of phage lysis. Molecular Microbiology, 21(4):675–682, 1996.

[82] Ry Young, Ing-Nang Wang, and William D. Roof. Phages will out: strategies of host cell lysis. Trends in Microbiology, 8(3):120–128, 2000.

[83] Abhyudai Singh and John J Dennehy. Stochastic holin expression can account for lysis time variation in the bacteriophage A. Journal of The Royal Society Interface, 11(95):20140140, 2014.

[84] Xiaoxin Liao, LQ Wang, and Pei Yu. Stability of Dynamical Systems, volume 5. Elsevier, 2007.

[85] Gurudas Chatterjee. Negative integral powers of a bidiagonal matrix. Mathematics of Computation, 28:713–714, 1974.

